# Rethinking cancer drug synergy prediction: a call for standardization in machine learning applications

**DOI:** 10.1101/2024.12.24.630216

**Authors:** Alexandra M. Wong, Lorin Crawford

**Affiliations:** Brown University Providence, RI 02912; Microsoft Research Cambridge, MA 02142

**Keywords:** cancer combination therapies *·* synergy prediction *·* machine learning *·* multi-omics

## Abstract

Drug resistance poses a significant challenge to cancer treatment, often caused by intratumor heterogeneity. Combination therapies have been shown to be an effective strategy to prevent resistant cancer cells from escaping single-drug treatments. However, discovering new drug combinations through traditional molecular assays can be costly and time-consuming. *In silico* approaches can overcome this limitation by exploring many candidate combinations at scale. This study systematically evaluates the utility of various machine learning algorithms, input features, and drug synergy prediction tasks. Our findings indicate a pressing need for establishing a standardized framework to measure and develop algorithms capable of predicting synergy.

## 1 Introduction

Cancer is the second leading cause of death in the United States [1, 2], and drug resistance is responsible for approximately 90% of deaths in patients receiving chemotherapy or targeted therapy. While drug resistance is caused by many mechanisms, tumor heterogeneity, where subpopulations of cells proliferate despite therapeutic pressure [3, 4], is one of the most significant mechanisms [5, 6]. Combination therapies have been proposed as an alternative strategy to overcome this limitation [7, 8]. Unfortunately, discovery of novel, patient-specific drug combinations is difficult due to costly molecular assay screening experiments [9, 10]. Large-scale genomic atlases and drug databases [11–13] have enabled the use of computational methods to systematically prioritize candidate combination therapies. However, there are differing approaches to performing drug synergy prediction, where researchers choose from multiple prediction tasks, input feature configurations, and machine learning algorithms.

First, most studies report one of three common tasks that are used to determine whether two drugs are synergistic when tested on a cell line: (1) binary classification of a synergy score, (2) synergy score regression, and (3) dose-dependent percent growth regression. The first prediction task binarizes numerical synergy scores, simply asking their models to classify whether pairs of drugs are synergistic at all on a given cell line [14–17]. Other methods perform regression on a continuous synergy score, detailing the degree to which two drugs will be synergistic [18–23]. Most of these studies overlook the importance of dosage information, a critical factor for both preclinical and clinical researchers. Synergy is often concentration dependent: for example, certain drug combinations can be synergistic at low concentrations but become antagonistic at high concentrations [24, 25]. To our knowledge, only ComboFM [26] and ComboPath [27] use the third prediction task where they predict the dose-dependent percent growth response of the cell line to a tested drug pair. This raises the question of whether optimizing for specific types of prediction tasks results in computational models that are more effective at selecting real drug combination candidates in practice.

Second, there is little consensus on the optimal set of input features needed to perform synergy prediction well. For example, some studies use Morgan fingerprints (MF), a binary vector representation of drug structure, combined with cell line DNA information [28]. Others use drug structure and RNA information instead [17, 29]. Many focus on the potential gains from a multi-omic approach with MF, leveraging information shared between DNA and RNA [19], combining RNA and protein expression [30], or utilizing DNA, RNA, and protein data altogether [31]. The inclusion of multi-modal features is often presumed to improve drug synergy predictions [14, 19, 23], so identifying the most predictive set of omic features could streamline future model development.

Lastly, there has yet to be a unified perspective on which computational methods are state-of-the-art for predicting combination therapies. Current approaches primarily focus on statistical and machine learning algorithms including random forests [32] and XGBoost [33]. However, more recent studies explore more complex frameworks. DeepSynergy [21], DrugCell [28], DeepMDS [34], MatchMaker [35], and DeepSignalingSynergy [36] use feed-forward deep neural networks with various multi-omic data as input features. Other researchers employ even deeper architectures [15, 17–19, 37, 38]. This motivates a systematic comparison of machine learning methods to determine their relative strengths and weaknesses in predicting drug synergy.

In this study, we conduct a systematic evaluation across synergy prediction tasks, multi-omic input feature combinations, and machine learning techniques. Each of our analyses lead to different conclusions about the best computational approach and highlight important inconsistencies in the field. Specifically, we find that reporting performance on just one type of prediction task is insufficient as different scores cover different aspects of what it means to be synergistic. Moreover, we find that incorporating multi-omic data does not automatically lead to improved model performance and that simple methods tend to perform just as well as those with complex architectures. We conclude our study with recommendations for the field of cancer drug synergy prediction.

## 2 Materials and Methods

### 2.1 Drug synergy scores

In this work, we use the NCI-ALMANAC database [11], which includes synergy scores for 3,686,475 drug combinations at varying doses to NCI-60 human tumor cell lines. We focus on performing three different tasks as it relates to predicting drug synergy: (1) binary classification of synergy, (2) synergy score regression, and (3) dose-dependent percent growth regression. The first task is to perform classification by binarizing the NCI ComboScore where (i) synergistic drug pairs are labeled as 1 and (ii) additive and antagonistic drug pairs are labeled as 0. The ComboScore, a modified version of the Bliss independence score, is used to determine the benefit of combining two drugs. Our second predictive task is to perform regression onto this continuous ComboScore.

Let *A* and *B* denote two drugs given at doses *p* and *q*, respectively. The ComboScore for a drug pair tested on a cell line is computed by summing over the differences between the expected (*Y_E_*) and observed (*Y_O_*) cell line percent growth to a drug combination treatment for all concentrations [11]:

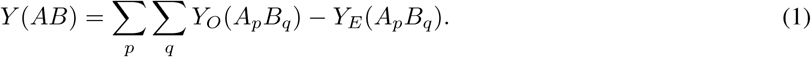

The expected percent growth for any combination *Y_E_*(*A_p_B_q_*) is based on the individual effect of each drug where [11]:

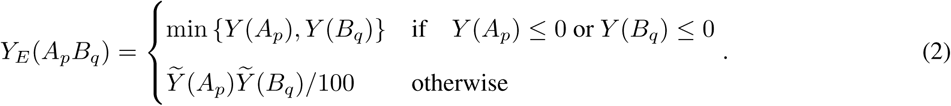

Intuitively, the expected combination growth falls into one of two cases. If one of the drugs *A* or *B* successfully kills the cancer entirely, then that value from the monotherapy is exactly the expected growth. Otherwise, the expected growth is the assumption that the two drugs would work independently and that their effects would multiply. Here, we follow previous work [11] and set *Ỹ* (*A_p_*) = min*{Y* (*A_p_*), 100*}* which truncates the effect of monotherapy for each drug at 100. This sets a 100% growth ceiling or, equivalently, that the expected combination percent growth should be less than the control. Thus, when a drug combination’s observed responses are greater than the expected percent growth responses (equivalent to a ComboScore greater than 0), that combination is considered synergistic [11].

The third synergy score we consider measures the percentage of cell growth (*Y_O_*(*A_p_B_q_*)) in the presence of a drug pair. The percent growth value is calculated at each drug concentration level using the following expression [39]:

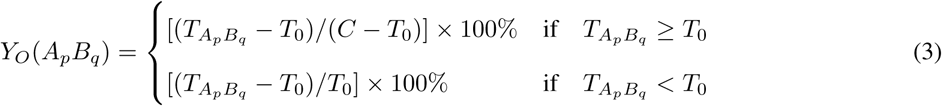

where *C* denotes control growth, *T*_0_ represents growth at time zero, and *T_Ap Bq_* is the test growth in presence of drug *A* at the concentration *p* and drug *B* at concentration *q*. The first of these cases calculates how much the drug combination successfully inhibited cell growth compared to the control. The second case assesses the percentage of cells that died when no additional growth occurred due to the treatment. Percent growth is a useful metric since it correlates with tumor growth and growth rate inhibition—both of which are better indicators of drug sensitivity to experimental researchers and clinicians in practice [40, 41]. By evaluating how machine learning methods perform dose-dependent prediction, we offer insight into their usefulness in the real world.

### 2.2 Machine learning models

We evaluate the ability of seven computational tools to predict drug synergy. Descriptions of how we implement these approaches are given below. The code for our analysis is publicly available (see Data and code availability).

#### Support vector machine

We implement a linear support vector machine (SVM) using the scikit-learn Python library [42]. To classify the binarized ComboScore synergy values, we use the LinearSVC flag with an *L*_2_-penalty, squared hinge loss, regularization parameter set to *λ* = 1, stopping criteria tolerance set to 0.0001, and 1000 maximum iterations. For regression-based tasks with the continuous ComboScore and dose-dependent percent growth values, we use the LinearSVR flag while setting the regularization parameter to *λ* = 1, the stopping criteria tolerance to 0.0001, and the maximum number of iterations to 1000. The LinearSVR flag implements an epsilon-insensitive loss function where we set *ɛ* = 0.

#### Gradient boosted decision trees

We implement a GPU compatible version of the “eXtreme Gradient Boosting” (XGBoost) algorithm from a Python package created by Chen and Guestrin [43]. For the binary classification task, we implement the XGBClassifier with the binary:hinge flag to make predictions of 0 (non-synergistic) or 1 (synergistic), rather than produce probabilities. All default parameters for the software are used, with the exception of setting the tree method to auto for scalability, the number of estimators (n_estimators) to 100, the maximum depth to 7, and the learning rate to 0.1. All the same parameters are used for the regression-based prediction tasks, where we implement the XGBRegressor model from the same Python package. The only differing parameter is the objective function, which is changed to reg:squarederror to minimize the squared error from the true ComboScore value.

#### Random forest

To implement the random forest (RF), we again turn to scikit-learn [42]. Here, we similarly use the RandomForestClassifier tag for classification of the binary ComboScore, and we use RandomForestRegressor for regression on the continuous ComboScore and percent growth values^1^. In each setting, we use 512 estimators (i.e., the number of trees) and keep the rest of the parameters to their default settings. For the classifier, this includes using a Gini criterion to measure the quality of a split in the tree; while a squared error loss is used for the regression models.

#### SYNDEEP

To compare against deep learning models, we also implement the SYNDEEP model by Torkamannia et al. [14]. This approach is considered “state-of-the-art” for binary classification of drug synergy on the NCI-ALMANAC data. Our PyTorch implementation contains a separate model for each of the binary classification and regression tasks. SYNDEEP has three hidden layers with 512, 128, and 32 nodes, respectively. For training, we use a 0.8 dropout rate, a 0.0002 learning rate, an Adam optimizer, and 300 epochs. Binary classification uses the binary cross entropy loss function while regression models use mean squared error loss.

#### Shallow and fully connected feedforward neural networks

We systematically evaluate multiple architectures of feedforward neural networks. The first model we refer to as a “shallow” neural network (SNN) consisting of a single hidden layer with 256 nodes. Here, we use an Adam optimizer with a 0.8 dropout rate and a 0.0002 learning rate for training. The parameters are chosen to facilitate comparison with SYNDEEP [14] (detailed in the previous section). We execute binary classification using the binary cross entropy loss function and regression-based models using the mean squared error loss function. We train all models for 300 epochs.

#### Partially connected neural networks

Recent research has developed customized neural network architectures that are inspired by biological systems [44]. Here, rather than fully connected and (often times) overparameterized models, these methods have partially connected architectures that are based on biological annotations in the literature or are derived from other functional relationships that have been identified through experimental validation. To explore how useful this type of architecture is for drug synergy prediction, we implement a partially connected neural network with a hidden gene layer (PCNNGL). This model contains a single layer where input DNA, RNA, and protein features are grouped together only if they correspond to the same gene (Supplementary Figure 1). MFs and drug concentration information are connected to each hidden node. We did not use dropout for the PCNNGL models since they already have sparse architectures. During training, we use 300 epochs, a learning rate of 0.0002, and an Adam optimizer for a similar comparison to SYNDEEP. Binary classification uses the binary cross entropy loss function while regression models use mean squared error loss.

#### PCNNGL-SYNDEEP

Lastly, we test a method that combines the architectures of PCNNGL and SYNDEEP. This hybrid model replaces the first hidden layer in SYNDEEP with the partially connected gene layer from PCNNGL, keeping the rest of the SYNDEEP model consistent. This results in a three hidden layer neural network with 722, 128, and 32 nodes, respectively. For consistency, we use an Adam optimizer, a 0.8 dropout rate, a 0.0002 learning rate, and 300 epochs for training. Binary classification uses the binary cross entropy loss function while regression models use mean squared error loss.

### 2.3 Data preprocessing

The NCI-ALMANAC contains 106 unique drugs and 61 cell lines representing nine distinct cancer subtypes. Whole exome DNA sequencing, gene expression via RNA sequencing, and protein expression data for NCI-60 cell lines were downloaded from CellMiner [45]. Drug features in the NCI-ALMANAC are encoded using the “Simplified

Molecular Input Line Entry System” (SMILES), a commonly adopted approach to convert chemical structures into string representations. For each drug, these SMILES strings are converted into 256-dimensional binary vectors called Morgan fingerprints (MF) using RDKit [46, 47]. This results in 512 drug structure features for each drug combination. Furthermore, in the percent growth prediction task, drug concentrations are also included as features in each machine learning model to account for dose dependence (bringing the total number of drug features to 514).

An outline of how the training data are constructed for each cell line can be found in Figure 1. Here, we only use cell lines that passed the quality control checks set by the NCI-ALMANAC. We also remove genomic features that have missing values. Originally, this leaves each cell line with a feature vector containing 23,372 DNA, 2665 RNA, and 2668 protein measurements. However, to maintain a balanced representation of modalities, we further limit the DNA data to the top 5% most variable features. After preprocessing, our final data set consists of 58 cell lines each with 1171 DNA, 722 RNA, and 722 protein features.

**Figure 1:**
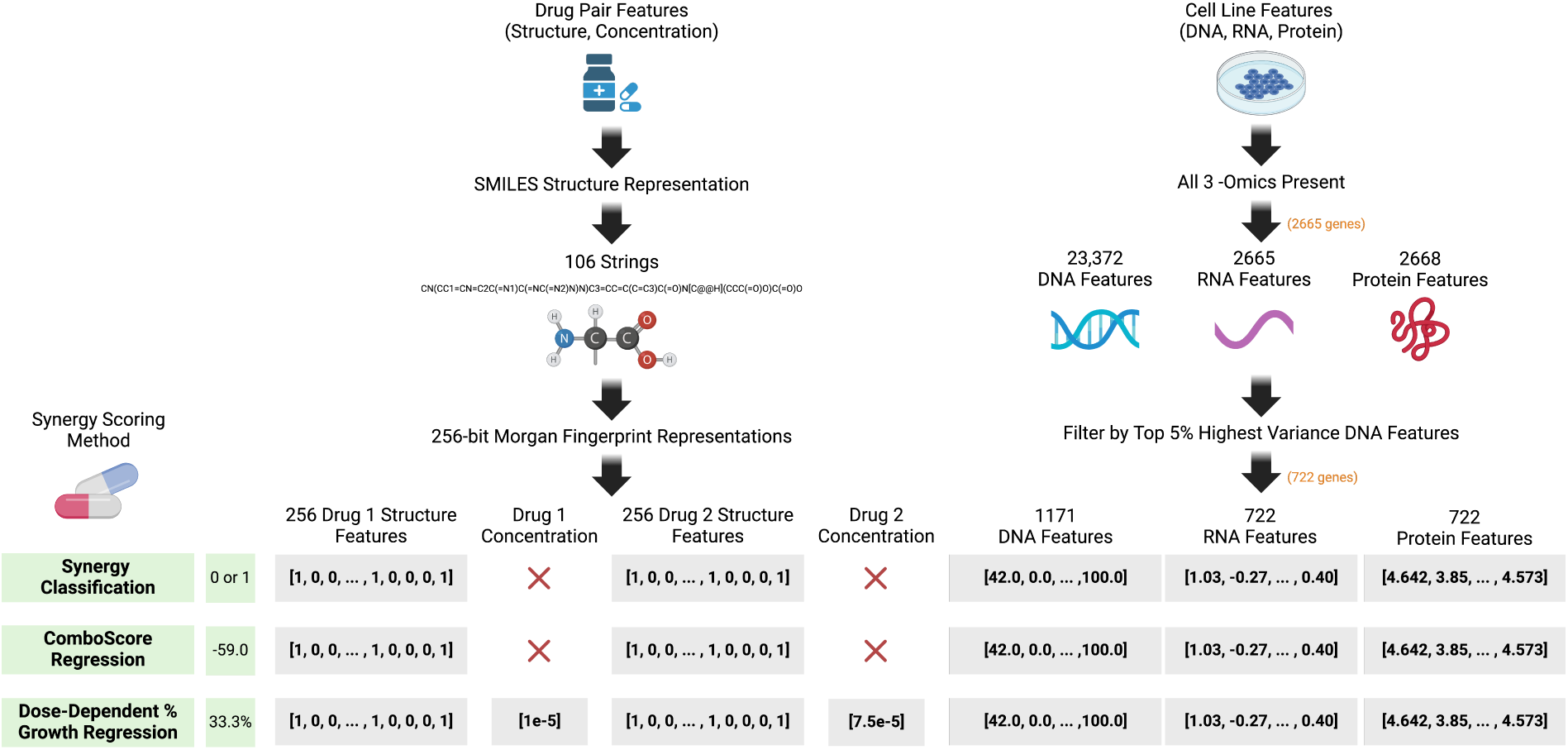
Schematic showing how feature vectors for each drug pair and NCI-60 cell line are constructed. The three types of synergy scores are shown in the bottom left corner as three green boxes with examples of each to the right. Synergy classification corresponds to the task of predicting binary labels 0 or 1 where 1 denotes a synergistic drug pair. ComboScore regression refers to the task of predicting the continuous ComboScore metric. Dose-dependent percent growth regression corresponds to predicting the expected percent growth of a cancer cell line in the presence of a particular drug combination at specific drug concentrations. The gray boxes show examples of the input features that are used for each prediction task. These features fall into one of two categories corresponding to either a drug or cell line (omic) feature. The left column of arrows displays the preprocessing of the drug features, while the right column of arrows shows the preprocessing of cell line features.

We perform 10-fold cross-validation for each analysis using the KFold function in scikit-learn. For the binary classification task, we enforce a balanced training subset where 50% of the observed drug pairs are synergistic and 50% are not.

## 3 Results

We evaluate the performance of machine learning algorithms at predicting drug synergy across different (i) types of scores, (ii) multi-omics input feature configurations, and (iii) model architectures. We report the average test accuracy for the binary classification task and the average Pearson correlation coefficient for the two regression tasks via a heatmap in Figure 2. Within each panel, results corresponding to each input feature configuration are given across the columns. These correspond to modeling Morgan fingerprints alone (MF), MF combined with a single modality (MF+DNA, MF+RNA, and MF+PE where PE represents protein expression data), MF combined with multiple omics (MF+DNA+RNA, MF+DNA+PE, MF+RNA+PE), and MF with all modalities (All). The test sensitivity, specificity, precision, F1 score, Matthew’s correlation coefficient (MCC), area under the curve (AUC), and the kappa statistic are also reported for the ComboScore classification task in Supplementary Table 3. For the ComboScore regression and percent growth regression tasks, we include the mean squared error (MSE), root mean squared error (RMSE), mean absolute error (MAE), the coefficient of determination (*R*^2^), and Spearman’s rank correlation in Supplementary Tables 4 and 5.

**Figure 2:**
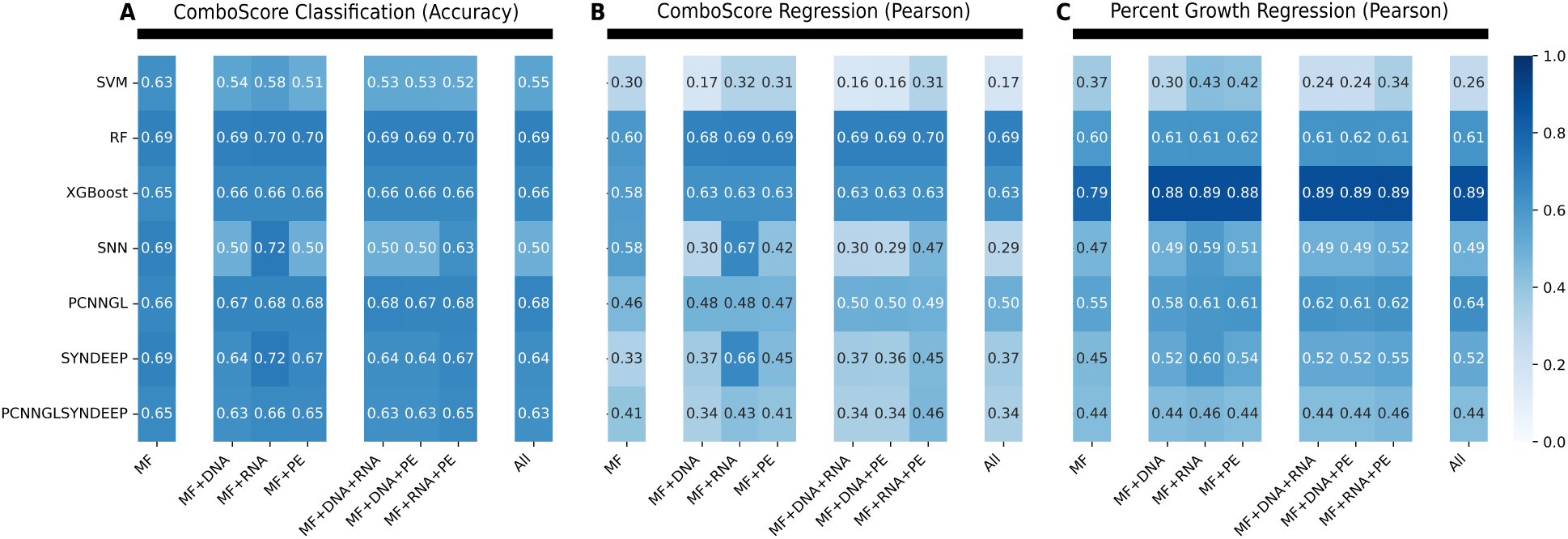
Drug synergy prediction results across different combinations of machine learning algorithms and omics measurements as input features. The *x*-axis displays which feature combination is used as input features for each model. The *y*-axis contains the labels for each machine learning algorithm. Results are averaged over the 10-fold cross-validation. **(A)** The average accuracy for each model and input feature combination on the binarized ComboScore synergy classification task. **(B)** Pearson correlation coefficient on the ComboScore regression task. **(C)** Pearson correlation coefficient on the percent growth regression task.

### 3.1 Binary ComboScore classification is a limited prediction benchmark

#### Performance between machine learning algorithms is indistinguishable when predicting binary synergy scores

There is very little variability in performance across the different machine learning algorithms for the classification task (Figure 2A). Out of the 56 method and input feature configurations, 42 (or 75%) result in accuracies that fall within the range of 0.60 to 0.70. While random forest has the highest accuracy in seven out of the eight input feature combinations, the difference between its accuracy and the second best model is negligible, only ranging from 1 *×* 10*^−^*^3^ to 0.02. While it is possible that this is simply the performance limit for the classification task on these data, this raises concerns about the discriminative power when predicting binary scores summarizing drug synergy. When different algorithmic architectures achieve similar results, it becomes challenging to meaningfully distinguish superior methodological approaches.

#### Success in binary classification does not correlate with performance in the regression task

We observe that a model’s ability to accurately classify the binary ComboScores does not necessarily translate to other synergy prediction challenges, particularly regression tasks. For example, the most accurate models in the classification task are SNN and SYNDEEP, which each score 0.72 in accuracy with the MF+RNA input feature combination. However, when examining these models in the regression tasks, neither has the highest Pearson correlation regardless of input data (Figure 2B and 2C). Instead, RF is the best method with the highest Pearson correlation at 0.70, contrasting with the best performance for both SNN (0.67) and SYNDEEP (0.66). Even when limiting the scope to the MF+RNA analysis in ComboScore regression, the Pearson correlation for RF at 0.69 still bests both SNN and SYNDEEP. In dose-dependent percent growth regression (Figure 2C), the top performing RF (0.62), XGBoost (0.89) and PCNNGL (0.61) models all show greater values than that of both SNN (0.59) and SYNDEEP (0.60).

This task-dependent performance is evident when examining XGBoost, which demonstrates an overall performance comparable to other models’ accuracies in the classification task, regardless of input features (Figure 2A). However, XGBoost’s average Pearson correlation (0.62) makes it the second best model when predicting the continuous ComboScore values and then the best performing model (0.88) in predicting percent growth (Figure 2C). This is notable when considering that, unlike some of the other models we chose to evaluate, XGBoost was not specifically designed for cancer drug synergy prediction. For example, we decided to analyze SYNDEEP because of its previously demonstrated high classification accuracy of drug pairs [14] and, indeed, it emerges as the top classifier in our study as well. However, focusing only on classification could falsely suggest that it is the best for synergy prediction. Models that excel in classifying synergistic combinations may falter in predicting exact synergy scores, while other algorithms perform better with different scoring metrics. Our results indicate that binary classification does not capture the full complexity of the drug synergy prediction problem.

#### Continuous synergy metrics lead to clear differentiating performances across predictive models

Assessing model performance in both regression tasks offers a clearer picture into the predictive power of different machine learning models than simple classification accuracy. Models using just MF alone to predict ComboScore (Figure 2B) and percent growth (Figure 2C) resulted in Pearson correlations ranging from of 0.30 to 0.79. This is in stark contrast to the classification task where models trained with MF only differed by 0.06 in terms of accuracy. A wider range allows for a clearer determination that RF and XGBoost are the top performing models in the two regression tasks. In ComboScore regression, the average RF Pearson correlation across modalities was 0.68, slightly outperforming XGBoost with an average of 0.62.

#### More complex models do not imply better performance

Our data in the regression tasks also allow for clearer comparison between models with varying complexity. For example, PCNNGL is a single hidden layer network while PCNNGL-SYNDEEP adds two additional layers from the SYNDEEP architecture. In both regression tasks, the PCNNGL-SYNDEEP model consistently performs worse than PCNNGL alone. Additionally, RF and XGBoost place as the top two models in the regression tasks, highlighting that deeper model architectures are not always guaranteed to outperform less complex methods.

### 3.2 Multi-omics data does not lead to model performance gains

We observe that adding more input data modalities does not necessarily improve model performance. In binary classification, combining MF with RNA expression often leads to one of the highest accuracy within each model (Figure 2A). However, the largest difference in accuracy between using MF+RNA as input features versus MF alone in any of these models is only 0.03. XGBoost records an average accuracy of 0.66 in all but the MF-only model, where it has a predictive accuracy of 0.65. Similarly, the predictive performance for each of the RF models only varies between 0.69 and 0.70 across all feature combinations.

We see a similar story in both ComboScore and percent growth regression tasks. Drug structure combined with gene expression (MF+RNA) is a top performing modality for SVM, XGBoost, SNN, and SYNDEEP. However, only SYNDEEP shows a notable performance increase compared to drug structure alone (MF). Additional omics features show no increase in benefit to model performance. In fact, combinations like MF+DNA+RNA, MF+RNA+PE, and MF+All (where all other data modalities are added to MF+RNA) have the ability to decrease the quality of many models. This is especially pronounced in SNN and SYNDEEP, which sees reductions in both accuracy and Pearson correlation coefficients in every prediction task (Figure 2). Taken together, this challenges the assumption that simply more data and additional features translates to better machine learning performance.

### 3.3 Model performance is robust across tissue type and drug pair classes

Given imbalanced sample sizes of cancer types and drug classes, one question that arises is whether evaluating machine learning methods on a multi-cancer dataset is appropriate. The NCI-ALMANAC dataset consists of six drug class categories and nine types of cancer: breast, central nervous system (CNS), colon, leukemia, melanoma, non-small cell lung carcinoma (NSCLC), ovarian, prostate, and renal) [11]. Therefore, we examine whether any model’s performance on specific tissue types or drug classes significantly differs from the results observed when analyzing all cancer types collectively.

In the ComboScore prediction tasks, the largest cancer type consists of 46,771 drug combinations for NSCLC. The tissue type with the least number of entries is the prostate with 10,541. In the percent growth regression scenario, the largest tissue type is also NSCLC with 430,041 entries, and the smallest is prostate at 97,191. For each cancer type, we performed a 10-fold cross-validation training and test scheme within each subset. For this analysis, we focus on XGBoost because it has the highest Pearson correlation coefficient values in the most clinically relevant prediction task: percent growth regression. We also focus our analysis to the drug structure and gene expression data combination, since it is the highest performing input feature set. Subclass performance for RF, SNN, PCNNGL, and SYNDEEP can be found in Supplementary Figures 2-9.

In all three prediction tasks, there is little variation in XGBoost performance between cancer types, showing that model performance robust to tissue subclasses with varying sample sizes (Figure 3). The tissue type results are also within range of the XGBoost performance on the whole dataset. This finding is consistent among all other models.

**Figure 3:**
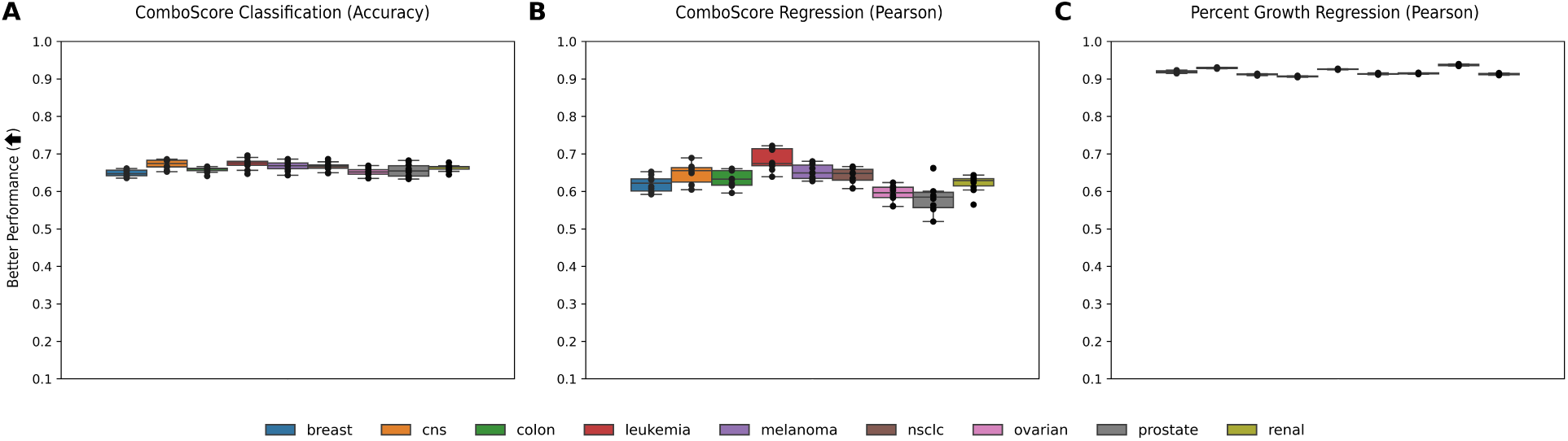
XGBoost performance on predicting different drug synergy scores across specific tissue types when trained on Morgan fingerprints (MF) and RNA expression data. **(A)** Synergy classification prediction task using accuracy as the primary metric. **(B)** ComboScore regression prediction task results using Pearson correlation coefficients. **(C)** Percent growth regression results using Pearson correlation coefficients. Results are based on 10-fold cross-validation. The box plots are drawn where the middle horizontal line corresponds to the median. The upper bound of the box denotes the 75th percentile while the lower bound marks the 25th percentile. The whiskers show the range of the distribution, and any points that are outside of the whiskers are determined to be outliers outside of 1.5 times the inter-quartile range (25th to 75th percentiles).

We also explore model robustness to specific drug classes using the same setup as in the cancer type breakdown. We show XGBoost’s performance on all prediction tasks using the MF+RNA data in Figure 4 using the same cross-validation scheme as in the cancer type breakdown. XGBoost is stable in all three prediction tasks, with an exception of the other_other category (neither drug belongs to the chemotherapy or targeted therapy classes) in the ComboScore regression task in Figure 4B. This sensitivity of ComboScore regression is likely explained by the drastic change in sample size for training: other_other is the smallest category containing only 8,814 samples. This is in contrast to the chemotherapy combination group (chemo_chemo) which is the largest subclass of ComboScores at 109,287 combinations. This lower performance is also reflected in the RF and SYNDEEP results (Supplementary Figures 7 and 9, respectively). This sample size difference does not significantly impact the performance of the percent growth regression task: the smallest category (other_other) at 79,326 samples maintains similar results to all other subclasses (Figure 4C). In general, model performance does not appear to be significantly different on particular subclasses of cancer types or drug classes versus analyzing the whole dataset together.

**Figure 4:**
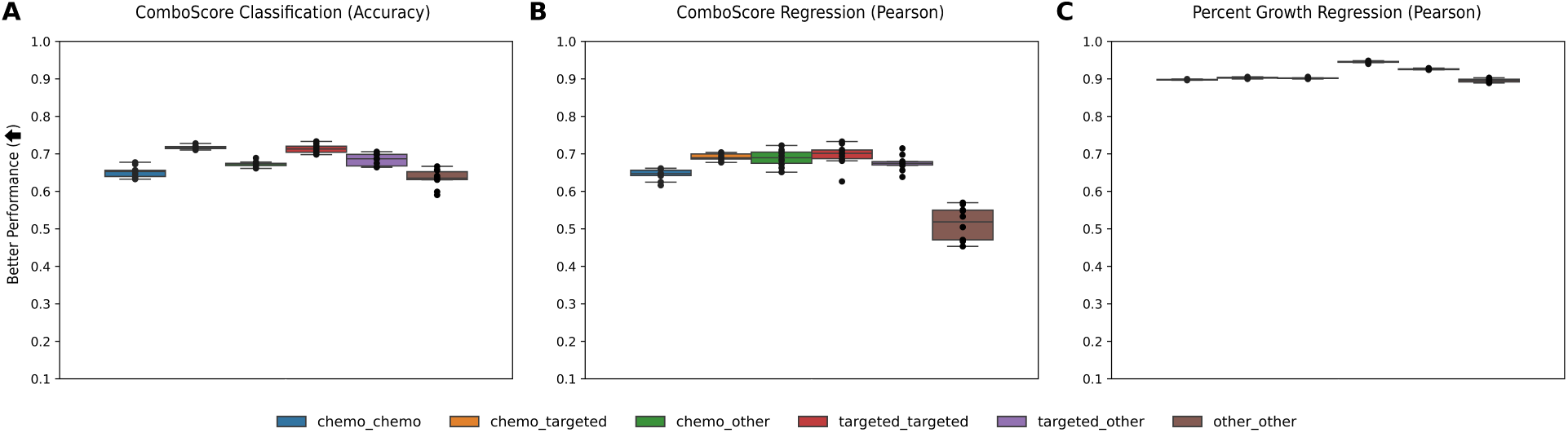
XGBoost performance on predicting different synergy scores across drug pair categories when trained on Morgan fingerprints (MF) and RNA expression data. Drugs are classified into one of three categories: chemotherapy, targeted therapy, or other. Drug pairs where both drugs are classified as chemotherapy are categorized as chemo_chemo. If one drug is a chemotherapy and the second drug is a targeted therapy, it is labeled as chemo_targeted. The remaining four categories of chemo_other, targeted_targeted, targeted_other, and other_other follow. **(A)** Synergy classification prediction task using accuracy as the primary metric. **(B)** ComboScore regression prediction task results using Pearson correlation coefficients. **(C)** Percent growth regression results using Pearson correlation coefficients. Results are based on 10-fold cross-validation. The box plots are drawn where the middle horizontal line corresponds to the median. The upper bound of the box denotes the 75th percentile while the lower bound marks the 25th percentile. The whiskers show the range of the distribution, and any points that are outside of the whiskers are determined to be outliers outside of 1.5 times the inter-quartile range (25th to 75th percentiles).

## 4 Discussion

In this study, we present a comprehensive evaluation of machine learning in cancer combination therapy prediction. Testing on the standardized NCI-ALMANAC dataset, we investigate three different ways to predict synergy, using eight unique input feature combinations across seven types of machine learning models. Our findings identify a critical need for standardization and challenge multi-omics and model complexity assumptions in the field of cancer drug synergy prediction.

First, our results indicate the importance of defining how drug synergy prediction is measured. The two main prediction tasks in the field include binary classification and synergy score regression. Our final scenario, dose-dependent percent growth regression, is less frequently used. The classification task accuracy is one of the most common versions of synergy prediction, yet our analyses found that it stratifies performance across machine learning model types and data combinations the least. Our findings show better performance stratification in both regression tasks, but we also find that stronger performance in one prediction task is not a signal of success in another.

There is still a wider debate on how to define binarized synergy. For classification, our study and Torkamannia et al. [14] use a cutoff of 0 to determine whether a given drug combination is considered synergistic. However, Wang et al. [17] use a much stricter cutoff to prioritize stronger combinations. There is also evidence that many clinically approved combination therapies exhibit primarily additive effects [48]. Therefore, the binary classification task of “greater-than-additive” synergy may be an oversimplification.

Our analyses indicate a crucial need for the field to standardize how synergy is predicted to appropriately compare methodologies. For *in silico* approaches to guide clinical cancer drug combination development, the field ought to align to metrics closer to the measurements that experimental researchers use (like dose-dependent percent growth regression). This not only helps researchers see differences in model behavior, but also maps more closely to the changes in tumor volume that experimental researchers and clinicians record. Percent growth regression also provides for greater precision in determining the dose-response curves, as ComboScores and other synergy scoring methods summarize over all concentrations. We recommend that all future studies report multiple synergy prediction tasks when assessing new machine learning models or data combinations, given the differences in relative model performance between prediction tasks.

Second, our results challenge the assumptions that “more data is better” and that “deeper models are better”. While our study observes that drug structure and gene expression data are often the stronger modality combination, our results also show that there is no significant advantage to a multi-omics strategy. This suggests that experimental researchers and clinicians may not gain new information when pursuing other omics modalities: combinations that added DNA or protein to the MF+RNA dataset often introduced more noise than MF+RNA alone. These data indicate that future studies should carefully evaluate whether additional modalities provide real performance gains. Furthermore, our results challenge the field’s tendency to produce more complex deep learning methods. RF and XGBoost models outperformed more sophisticated methods in all but one analysis, suggesting that RF and XGBoost should be standard benchmarks for all future synergy prediction studies. Our analysis also demonstrates that model performance on specific cancer types and drug classes is generally equivalent to the performance in the overall multi-cancer dataset.

This study has several limitations that should be addressed in future research. The analysis relies solely on Holbeck et al. [11]’s NCI-ALMANAC dataset which, while extensive, could be complemented by incorporating additional databases such as DrugComb ([13]) and O’Neil [12] to enhance the generalizability of the findings. There are also additional synergy scoring methods beyond the percent growth measure and ComboScores (a modified Bliss independence score), including highest single agent scores, zero interaction potency scores, and Loewe scores [13]. The field should investigate and choose standard metrics for reporting synergy to aid future comparisons. Additionally, the feature space in our study is confined to Morgan fingerprints, drug concentration, and multi-omics data (DNA, RNA, and protein). Future modalities for evaluation could include metabolomics, drug-target interaction networks, cellular signaling pathway information, and other physicochemical properties of the compounds. This study’s examination of seven different machine learning approaches should also be expanded to include numerous emerging modeling techniques in the field (e.g., those related to artificial intelligence). Such extensions could provide richer insights into drug synergy prediction and potentially improve prediction accuracy.

In conclusion, cancer drug synergy prediction is not only a combinatorially explosive space *in vitro* and *in vivo*, but also *in silico*. There are numerous ways to quantify synergy, modalities that can be used as predictive features, and computational algorithms from which to choose. Our study displays a crucial need for standardization when predicting synergy. The insights provided in this study have the potential to streamline synergy prediction efforts, enabling more efficient exploration of therapeutic combinations and guiding future research toward the most promising approaches. By establishing a clearer direction, we hope this work contributes to the advancement of the discovery and personalization of combination therapy strategies for cancer moving forward.

## Acknowledgments

We gratefully acknowledge Barbara Engelhardt (Stanford University), Sean Lawler (Brown University), and Ying Ma (Brown University) for their valuable feedback and insightful discussions that helped shape this work. We also thank Helen Xie (Brown University) for her assistance in background research that provided useful context for this study, and Alan DenAdel (Brown University) and Julian Stamp (Brown University) for useful comments on early drafts of the manuscript. This research was conducted in part using computational resources and services at the Center for Computation and Visualization at Brown University.

## Data and code availability

All of the data in this study is derived from publicly available sources. The NCI-ALMANAC dataset is available at https://wiki.nci.nih.gov/display/NCIDTPdata/NCI-ALMANAC. The multi-modal omics data are downloaded from CellMiner: DNA exome sequencing data can be found at https://discover.nci.nih.gov/cellminer/download/processeddataset/nci60_DNA Exome_Seq_none.zip, the RNA expression data at https://discov er.nci.nih.gov/cellminer/download/processeddataset/nci60_RNA 5_Platform_Gene_Transcript_Average_z_scores.zip, and the protein expression data at https://discover.nci.nih.gov/cellminer/dow nload/processeddataset/nci60_Protein SWATH_(Mass_spectrometry)_Protein.zip. The code written for this project is publicly available at https://github.com/lcrawlab/cancer-drug-synergy-prediction.

## Author contributions

AMW and LC conceived the study. AMW wrote the code and performed the analysis. LC supervised the project and provided resources. All authors wrote and revised the manuscript.

## Supplementary Material

**Supplemental Table 1:** Average Random Forest (RF) performance on the ComboScore classification task. The first section corresponds to the average across 10-fold cross validation splits using the CPU implementation of RF in scikit-learn. The second section contains the average across 10-fold cross validation splits using the GPU implementation of RF via XGBoost [43]. The vertical columns represent each of the input feature combinations: Morgan fingerprints (MF), MF with DNA, MF with RNA, MF with protein expression (PE), MF with DNA and RNA, MF with DNA and PE, MF with RNA and PE, and MF with all cell line feature modalities.

| CPU Implementation of RF |  |  |  |  |  |  |  |  |
| --- | --- | --- | --- | --- | --- | --- | --- | --- |
| Metric | MF | MF+DNA | MF+RNA | MF+PE | MF+DNA+RNA | MF+DNA+PE | MF+RNA+PE | MF+All |
| Accuracy | 0.688 | 0.695 | 0.702 | 0.702 | 0.693 | 0.693 | 0.695 | 0.688 |
| Sensitivity | 0.678 | 0.674 | 0.682 | 0.680 | 0.674 | 0.675 | 0.675 | 0.671 |
| Specificity | 0.697 | 0.715 | 0.723 | 0.723 | 0.712 | 0.712 | 0.715 | 0.705 |
| Precision | 0.692 | 0.703 | 0.711 | 0.711 | 0.700 | 0.701 | 0.704 | 0.695 |
| F1 Score | 0.684 | 0.688 | 0.696 | 0.695 | 0.687 | 0.688 | 0.689 | 0.683 |
| MCC | 0.376 | 0.390 | 0.405 | 0.404 | 0.386 | 0.387 | 0.391 | 0.376 |
| AUC | 0.688 | 0.695 | 0.702 | 0.702 | 0.693 | 0.693 | 0.695 | 0.688 |
| Kappa | 0.376 | 0.390 | 0.405 | 0.403 | 0.386 | 0.387 | 0.391 | 0.376 |
| GPU Implementation of RF |  |  |  |  |  |  |  |  |
| Metric | MF | MF+DNA | MF+RNA | MF+PE | MF+DNA+RNA | MF+DNA+PE | MF+RNA+PE | MF+All |
| Accuracy | 0.552 | 0.552 | 0.552 | 0.579 | 0.567 | 0.567 | 0.578 | 0.568 |
| Sensitivity | 0.222 | 0.226 | 0.231 | 0.346 | 0.289 | 0.284 | 0.341 | 0.290 |
| Specificity | 0.878 | 0.876 | 0.872 | 0.811 | 0.843 | 0.847 | 0.813 | 0.843 |
| Precision | 0.260 | 0.259 | 0.258 | 0.389 | 0.325 | 0.327 | 0.389 | 0.325 |
| F1 Score | 0.240 | 0.242 | 0.244 | 0.366 | 0.306 | 0.304 | 0.363 | 0.307 |
| MCC | 0.101 | 0.103 | 0.103 | 0.157 | 0.133 | 0.132 | 0.155 | 0.134 |
| AUC | 0.550 | 0.551 | 0.551 | 0.578 | 0.566 | 0.565 | 0.577 | 0.566 |
| Kappa | 0.100 | 0.102 | 0.103 | 0.156 | 0.132 | 0.131 | 0.154 | 0.133 |

**Supplemental Table 2:**
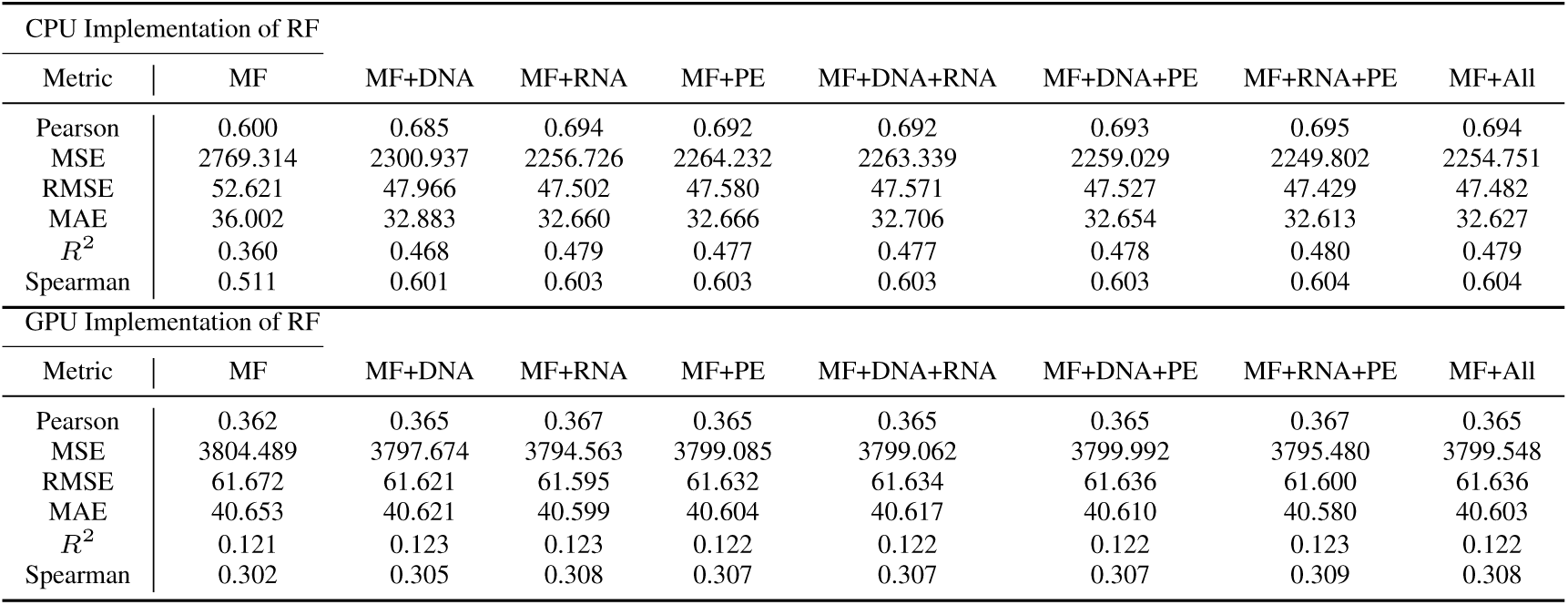
Average Random Forest (RF) performance on the ComboScore regression task. The first section corresponds to the average across 10-fold cross validation metrics using the CPU implementation of RF in scikit-learn. The second section contains the average across 10-fold cross validation splits using the GPU implementation of RF via XGBoost [43]. The vertical columns represent each of the input feature combinations: Morgan fingerprints (MF), MF with DNA, MF with RNA, MF with protein expression (PE), MF with DNA and RNA, MF with DNA and PE, MF with RNA and PE, and MF with all cell line feature modalities.

| CPU Implementation of RF |  |  |  |  |  |  |  |  |
| --- | --- | --- | --- | --- | --- | --- | --- | --- |
| Metric | MF | MF+DNA | MF+RNA | MF+PE | MF+DNA+RNA | MF+DNA+PE | MF+RNA+PE | MF+All |
| Pearson | 0.600 | 0.685 | 0.694 | 0.692 | 0.692 | 0.693 | 0.695 | 0.694 |
| MSE | 2769.314 | 2300.937 | 2256.726 | 2264.232 | 2263.339 | 2259.029 | 2249.802 | 2254.751 |
| RMSE | 52.621 | 47.966 | 47.502 | 47.580 | 47.571 | 47.527 | 47.429 | 47.482 |
| MAE | 36.002 | 32.883 | 32.660 | 32.666 | 32.706 | 32.654 | 32.613 | 32.627 |
| $R^2$ | 0.360 | 0.468 | 0.479 | 0.477 | 0.477 | 0.478 | 0.480 | 0.479 |
| Spearman | 0.511 | 0.601 | 0.603 | 0.603 | 0.603 | 0.603 | 0.604 | 0.604 |
| GPU Implementation of RF |  |  |  |  |  |  |  |  |
| Metric | MF | MF+DNA | MF+RNA | MF+PE | MF+DNA+RNA | MF+DNA+PE | MF+RNA+PE | MF+All |
| Pearson | 0.362 | 0.365 | 0.367 | 0.365 | 0.365 | 0.365 | 0.367 | 0.365 |
| MSE | 3804.489 | 3797.674 | 3794.563 | 3799.085 | 3799.062 | 3799.992 | 3795.480 | 3799.548 |
| RMSE | 61.672 | 61.621 | 61.595 | 61.632 | 61.634 | 61.636 | 61.600 | 61.636 |
| MAE | 40.653 | 40.621 | 40.599 | 40.604 | 40.617 | 40.610 | 40.580 | 40.603 |
| $R^2$ | 0.121 | 0.123 | 0.123 | 0.122 | 0.122 | 0.122 | 0.123 | 0.122 |
| Spearman | 0.302 | 0.305 | 0.308 | 0.307 | 0.307 | 0.307 | 0.309 | 0.308 |

**Supplemental Table 3:** Mean and standard deviation across 10-fold cross validation performance for seven different machine learning models on the ComboScore classification task. Each section corresponds to one of the following machine learning models including: support vector machine (SVM), random forest, eXtreme gradient boosting (XGBoost), shallow neural network (SNN), SYNDEEP [14], partially connected neural network with hidden gene layer (PCNNGL), and the PCNNGL-SYNDEEP hybrid model. The columns denote different input feature combinations: Morgan fingerprints (MF), MF with DNA, MF with RNA, MF with protein expression (PE), MF with DNA and RNA, MF with DNA and PE, MF with RNA and PE, and MF with all cell line feature modalities. Metrics in bold indicate the best performance across all input feature combinations for that model.

| Metric | MF | MF+DNA | MF+RNA | MF+PE | MF+DNA+RNA | MF+DNA+PE | MF+RNA+PE | MF+All |
| --- | --- | --- | --- | --- | --- | --- | --- | --- |
| **SVM** |  |  |  |  |  |  |  |  |
| Accuracy | **0.632 ± 0.003** | 0.542 ± 0.019 | 0.578 ± 0.008 | 0.512 ± 0.019 | 0.535 ± 0.019 | 0.533 ± 0.024 | 0.524 ± 0.023 | 0.546 ± 0.013 |
| Sensitivity | 0.611 ± 0.011 | 0.453 ± 0.324 | 0.574 ± 0.076 | 0.511 ± 0.505 | 0.424 ± 0.369 | 0.376 ± 0.347 | **0.814 ± 0.357** | 0.491 ± 0.296 |
| Specificity | 0.653 ± 0.007 | 0.628 ± 0.311 | 0.582 ± 0.078 | 0.515 ± 0.508 | 0.647 ± 0.348 | **0.692 ± 0.348** | 0.235 ± 0.383 | 0.600 ± 0.292 |
| Precision | 0.638 ± 0.006 | 0.609 ± 0.106 | 0.580 ± 0.014 | **0.698 ± 0.205** | 0.594 ± 0.063 | 0.608 ± 0.075 | 0.563 ± 0.113 | 0.580 ± 0.055 |
| F1 Score | **0.624 ± 0.006** | 0.431 ± 0.224 | 0.574 ± 0.032 | 0.365 ± 0.323 | 0.391 ± 0.245 | 0.370 ± 0.223 | 0.583 ± 0.193 | 0.474 ± 0.172 |
| MCC | **0.265 ± 0.007** | 0.105 ± 0.024 | 0.158 ± 0.016 | 0.077 ± 0.056 | 0.099 ± 0.031 | 0.095 ± 0.033 | 0.102 ± 0.063 | 0.112 ± 0.015 |
| AUC | **0.632 ± 0.003** | 0.541 ± 0.019 | 0.578 ± 0.008 | 0.513 ± 0.017 | 0.536 ± 0.019 | 0.534 ± 0.023 | 0.525 ± 0.024 | 0.545 ± 0.012 |
| Kappa | **0.265 ± 0.007** | 0.081 ± 0.038 | 0.156 ± 0.017 | 0.026 ± 0.035 | 0.071 ± 0.037 | 0.068 ± 0.045 | 0.049 ± 0.048 | 0.091 ± 0.024 |
| **Random Forest** |  |  |  |  |  |  |  |  |
| Accuracy | 0.689 ± 0.002 | 0.695 ± 0.003 | **0.702 ± 0.003** | **0.702 ± 0.003** | 0.693 ± 0.004 | 0.693 ± 0.003 | 0.695 ± 0.004 | 0.688 ± 0.004 |
| Sensitivity | 0.674 ± 0.006 | 0.674 ± 0.005 | **0.682 ± 0.004** | 0.680 ± 0.006 | 0.674 ± 0.006 | 0.675 ± 0.003 | 0.675 ± 0.005 | 0.671 ± 0.006 |
| Specificity | 0.705 ± 0.006 | 0.715 ± 0.003 | **0.723 ± 0.007** | **0.723 ± 0.005** | 0.712 ± 0.004 | 0.712 ± 0.005 | 0.715 ± 0.004 | 0.705 ± 0.003 |
| Precision | 0.695 ± 0.005 | 0.703 ± 0.004 | **0.711 ± 0.006** | **0.711 ± 0.006** | 0.700 ± 0.007 | 0.701 ± 0.005 | 0.704 ± 0.005 | 0.695 ± 0.004 |
| F1 Score | 0.685 ± 0.003 | 0.688 ± 0.004 | **0.696 ± 0.003** | 0.695 ± 0.004 | 0.687 ± 0.006 | 0.688 ± 0.004 | 0.689 ± 0.004 | 0.683 ± 0.004 |
| MCC | 0.379 ± 0.004 | 0.390 ± 0.007 | **0.405 ± 0.006** | 0.404 ± 0.006 | 0.386 ± 0.008 | 0.387 ± 0.006 | 0.391 ± 0.008 | 0.376 ± 0.007 |
| AUC | 0.689 ± 0.002 | 0.695 ± 0.003 | **0.702 ± 0.003** | **0.702 ± 0.003** | 0.693 ± 0.004 | 0.693 ± 0.003 | 0.695 ± 0.004 | 0.688 ± 0.004 |
| Kappa | 0.379 ± 0.004 | 0.390 ± 0.007 | **0.405 ± 0.006** | 0.403 ± 0.006 | 0.386 ± 0.008 | 0.387 ± 0.006 | 0.391 ± 0.008 | 0.376 ± 0.007 |
| **XGBoost** |  |  |  |  |  |  |  |  |
| Accuracy | 0.654 ± 0.005 | 0.661 ± 0.005 | 0.657 ± 0.009 | 0.657 ± 0.008 | 0.656 ± 0.009 | 0.657 ± 0.008 | **0.660 ± 0.008** | 0.657 ± 0.007 |
| Sensitivity | 0.632 ± 0.063 | 0.667 ± 0.083 | 0.691 ± 0.083 | 0.696 ± 0.078 | 0.704 ± 0.084 | 0.689 ± 0.086 | 0.677 ± 0.081 | **0.713 ± 0.070** |
| Specificity | **0.676 ± 0.069** | 0.656 ± 0.088 | 0.624 ± 0.100 | 0.618 ± 0.092 | 0.610 ± 0.100 | 0.626 ± 0.101 | 0.643 ± 0.092 | 0.602 ± 0.082 |
| Precision | **0.664 ± 0.027** | **0.664 ± 0.032** | 0.653 ± 0.037 | 0.650 ± 0.033 | 0.649 ± 0.038 | 0.654 ± 0.037 | 0.659 ± 0.033 | 0.645 ± 0.031 |
| F1 Score | 0.645 ± 0.019 | 0.661 ± 0.024 | 0.667 ± 0.020 | 0.668 ± 0.019 | 0.670 ± 0.020 | 0.666 ± 0.022 | 0.664 ± 0.021 | **0.674 ± 0.018** |
| MCC | 0.311 ± 0.010 | **0.327 ± 0.007** | 0.321 ± 0.014 | 0.320 ± 0.013 | 0.320 ± 0.014 | 0.321 ± 0.013 | 0.324 ± 0.012 | 0.321 ± 0.012 |
| AUC | 0.654 ± 0.004 | **0.661 ± 0.004** | 0.658 ± 0.009 | 0.657 ± 0.008 | 0.657 ± 0.008 | 0.658 ± 0.008 | 0.660 ± 0.008 | 0.658 ± 0.007 |
| Kappa | 0.308 ± 0.009 | **0.323 ± 0.009** | 0.315 ± 0.018 | 0.314 ± 0.016 | 0.314 ± 0.017 | 0.315 ± 0.017 | 0.320 ± 0.015 | 0.316 ± 0.015 |
| **SNN** |  |  |  |  |  |  |  |  |
| Accuracy | 0.687 ± 0.003 | 0.497 ± 0.003 | **0.716 ± 0.004** | 0.496 ± 0.003 | 0.498 ± 0.003 | 0.499 ± 0.002 | 0.634 ± 0.006 | 0.497 ± 0.003 |
| Sensitivity | 0.657 ± 0.006 | 0.302 ± 0.482 | **0.691 ± 0.008** | 0.500 ± 0.527 | 0.400 ± 0.516 | 0.600 ± 0.516 | 0.508 ± 0.062 | 0.400 ± 0.516 |
| Specificity | 0.718 ± 0.005 | 0.698 ± 0.482 | **0.740 ± 0.006** | 0.500 ± 0.527 | 0.600 ± 0.516 | 0.400 ± 0.516 | 0.761 ± 0.054 | 0.600 ± 0.516 |
| Precision | 0.700 ± 0.005 | 0.202 ± 0.261 | **0.727 ± 0.005** | 0.248 ± 0.262 | 0.199 ± 0.257 | 0.299 ± 0.258 | 0.683 ± 0.021 | 0.199 ± 0.256 |
| F1 Score | 0.677 ± 0.004 | 0.203 ± 0.318 | **0.708 ± 0.005** | 0.332 ± 0.350 | 0.266 ± 0.343 | 0.400 ± 0.344 | 0.579 ± 0.032 | 0.265 ± 0.343 |
| MCC | 0.375 ± 0.006 | 0.001 ± 0.003 | **0.432 ± 0.008** | 0.000 ± 0.000 | 0.000 ± 0.000 | 0.000 ± 0.000 | 0.279 ± 0.007 | 0.000 ± 0.000 |
| AUC | 0.687 ± 0.003 | 0.500 ± 0.000 | **0.716 ± 0.004** | 0.500 ± 0.000 | 0.500 ± 0.000 | 0.500 ± 0.000 | 0.634 ± 0.005 | 0.500 ± 0.000 |
| Kappa | 0.375 ± 0.006 | 0.000 ± 0.001 | **0.431 ± 0.008** | 0.000 ± 0.000 | 0.000 ± 0.000 | 0.000 ± 0.000 | 0.268 ± 0.011 | 0.000 ± 0.000 |
| **SYNDEEP** |  |  |  |  |  |  |  |  |
| Accuracy | 0.687 ± 0.003 | 0.639 ± 0.004 | **0.716 ± 0.004** | 0.667 ± 0.004 | 0.637 ± 0.004 | 0.638 ± 0.003 | 0.668 ± 0.005 | 0.639 ± 0.004 |
| Sensitivity | 0.660 ± 0.024 | 0.639 ± 0.038 | **0.683 ± 0.024** | 0.634 ± 0.037 | 0.630 ± 0.051 | 0.628 ± 0.029 | 0.642 ± 0.037 | 0.630 ± 0.027 |
| Specificity | 0.715 ± 0.022 | 0.639 ± 0.038 | **0.750 ± 0.024** | 0.701 ± 0.041 | 0.643 ± 0.052 | 0.649 ± 0.028 | 0.693 ± 0.040 | 0.648 ± 0.025 |
| Precision | 0.699 ± 0.007 | 0.639 ± 0.012 | **0.733 ± 0.011** | 0.681 ± 0.021 | 0.640 ± 0.014 | 0.642 ± 0.009 | 0.678 ± 0.015 | 0.642 ± 0.011 |
| F1 Score | 0.678 ± 0.010 | 0.638 ± 0.015 | **0.706 ± 0.009** | 0.656 ± 0.011 | 0.634 ± 0.019 | 0.634 ± 0.011 | 0.659 ± 0.013 | 0.635 ± 0.009 |
| MCC | 0.376 ± 0.005 | 0.278 ± 0.009 | **0.434 ± 0.008** | 0.337 ± 0.009 | 0.275 ± 0.008 | 0.277 ± 0.006 | 0.337 ± 0.009 | 0.279 ± 0.007 |
| AUC | 0.687 ± 0.003 | 0.639 ± 0.004 | **0.716 ± 0.004** | 0.668 ± 0.004 | 0.637 ± 0.004 | 0.638 ± 0.003 | 0.668 ± 0.005 | 0.639 ± 0.004 |
| Kappa | 0.375 ± 0.006 | 0.277 ± 0.009 | **0.433 ± 0.008** | 0.335 ± 0.007 | 0.274 ± 0.008 | 0.277 ± 0.006 | 0.336 ± 0.009 | 0.278 ± 0.007 |
| **PCNNGL** |  |  |  |  |  |  |  |  |
| Accuracy | 0.664 ± 0.004 | 0.674 ± 0.005 | 0.675 ± 0.003 | 0.675 ± 0.004 | 0.676 ± 0.003 | 0.675 ± 0.003 | 0.675 ± 0.003 | **0.677 ± 0.004** |
| Sensitivity | 0.641 ± 0.012 | 0.650 ± 0.017 | 0.647 ± 0.015 | 0.654 ± 0.014 | 0.652 ± 0.009 | **0.656 ± 0.015** | 0.653 ± 0.013 | 0.652 ± 0.016 |
| Specificity | 0.688 ± 0.013 | 0.698 ± 0.019 | **0.703 ± 0.016** | 0.697 ± 0.016 | 0.700 ± 0.010 | 0.693 ± 0.015 | 0.698 ± 0.012 | 0.702 ± 0.013 |
| Precision | 0.673 ± 0.007 | 0.683 ± 0.009 | **0.686 ± 0.008** | 0.683 ± 0.009 | 0.685 ± 0.005 | 0.682 ± 0.007 | 0.684 ± 0.007 | **0.686 ± 0.006** |
| F1 Score | 0.656 ± 0.006 | 0.666 ± 0.007 | 0.666 ± 0.005 | 0.668 ± 0.005 | 0.668 ± 0.004 | **0.669 ± 0.006** | 0.668 ± 0.004 | 0.668 ± 0.007 |
| MCC | 0.329 ± 0.007 | 0.349 ± 0.009 | 0.351 ± 0.005 | 0.351 ± 0.009 | 0.352 ± 0.006 | 0.350 ± 0.006 | 0.351 ± 0.005 | **0.354 ± 0.007** |
| AUC | 0.664 ± 0.004 | 0.674 ± 0.005 | 0.675 ± 0.003 | 0.675 ± 0.004 | 0.676 ± 0.003 | 0.675 ± 0.003 | 0.675 ± 0.002 | **0.677 ± 0.004** |
| Kappa | 0.329 ± 0.007 | 0.349 ± 0.009 | 0.350 ± 0.005 | 0.351 ± 0.009 | 0.352 ± 0.006 | 0.350 ± 0.006 | 0.351 ± 0.005 | **0.354 ± 0.007** |
| **PCNNGL-SYNDEEP** |  |  |  |  |  |  |  |  |
| Accuracy | 0.653 ± 0.003 | 0.631 ± 0.004 | **0.656 ± 0.003** | 0.652 ± 0.004 | 0.631 ± 0.004 | 0.631 ± 0.004 | 0.655 ± 0.005 | 0.629 ± 0.005 |
| Sensitivity | 0.559 ± 0.030 | 0.543 ± 0.051 | **0.583 ± 0.023** | 0.567 ± 0.029 | 0.528 ± 0.041 | 0.533 ± 0.035 | 0.570 ± 0.031 | 0.520 ± 0.040 |
| Specificity | **0.747 ± 0.030** | 0.721 ± 0.046 | 0.729 ± 0.020 | 0.737 ± 0.026 | 0.733 ± 0.037 | 0.729 ± 0.031 | 0.739 ± 0.026 | 0.737 ± 0.035 |
| Precision | **0.689 ± 0.015** | 0.662 ± 0.017 | 0.683 ± 0.009 | 0.683 ± 0.013 | 0.665 ± 0.013 | 0.663 ± 0.012 | 0.687 ± 0.011 | 0.666 ± 0.012 |
| F1 Score | 0.617 ± 0.013 | 0.594 ± 0.024 | **0.628 ± 0.010** | 0.619 ± 0.013 | 0.588 ± 0.020 | 0.590 ± 0.018 | 0.622 ± 0.015 | 0.583 ± 0.022 |
| MCC | 0.312 ± 0.007 | 0.269 ± 0.005 | **0.315 ± 0.004** | 0.308 ± 0.008 | 0.268 ± 0.007 | 0.267 ± 0.008 | 0.314 ± 0.009 | 0.265 ± 0.009 |
| AUC | 0.653 ± 0.003 | 0.632 ± 0.004 | **0.656 ± 0.003** | 0.652 ± 0.004 | 0.631 ± 0.004 | 0.631 ± 0.005 | 0.655 ± 0.005 | 0.629 ± 0.005 |
| Kappa | 0.306 ± 0.006 | 0.263 ± 0.007 | **0.311 ± 0.005** | 0.304 ± 0.009 | 0.261 ± 0.008 | 0.262 ± 0.009 | 0.309 ± 0.011 | 0.258 ± 0.010 |

**Supplemental Table 4:** Mean and standard deviation across 10-fold cross validation performance for seven different machine learning models on the ComboScore regression task. Each section corresponds to one of the following machine learning models including: support vector machine (SVM), random forest, eXtreme gradient boosting (XGBoost), shallow neural network (SNN), SYNDEEP [14], partially connected neural network with hidden gene layer (PCNNGL), and the PCNNGL-SYNDEEP hybrid model. The columns denote different input feature combinations: Morgan fingerprints (MF), MF with DNA, MF with RNA, MF with protein expression (PE), MF with DNA and RNA, MF with DNA and PE, MF with RNA and PE, and MF with all cell line feature modalities. Metrics in bold indicate the best performance across all input feature combinations for that model.

| Metric | MF | MF+DNA | MF+RNA | MF+PE | MF+DNA+RNA | MF+DNA+PE | MF+RNA+PE | MF+All |
| --- | --- | --- | --- | --- | --- | --- | --- | --- |
| **SVM** |  |  |  |  |  |  |  |  |
| MSE | 3966.588 ± 155.840 | 5777.938 ± 517.024 | **3912.852 ± 106.891** | 3972.874 ± 106.386 | 9575.918 ± 5312.605 | 7242.081 ± 3209.592 | 4062.530 ± 220.633 | 6012.979 ± 1275.457 |
| RMSE | 62.970 ± 1.234 | 75.946 ± 3.364 | **62.548 ± 0.856** | 63.026 ± 0.837 | 94.752 ± 25.775 | 83.705 ± 16.180 | 63.716 ± 1.750 | 77.193 ± 7.757 |
| MAE | 40.531 ± 0.416 | 55.131 ± 3.689 | **40.165 ± 0.227** | 40.881 ± 0.782 | 74.249 ± 26.270 | 63.643 ± 17.043 | 41.494 ± 1.392 | 56.381 ± 8.525 |
| $R^2$ | 0.084 ± 0.003 | -0.336 ± 0.131 | **0.096 ± 0.003** | 0.082 ± 0.014 | -1.210 ± 1.229 | -0.676 ± 0.753 | 0.062 ± 0.036 | -0.389 ± 0.292 |
| Pearson | 0.296 ± 0.005 | 0.171 ± 0.021 | **0.316 ± 0.006** | 0.315 ± 0.005 | 0.157 ± 0.033 | 0.160 ± 0.014 | 0.313 ± 0.006 | 0.171 ± 0.017 |
| Spearman | 0.331 ± 0.005 | 0.182 ± 0.015 | **0.356 ± 0.004** | **0.356 ± 0.004** | 0.162 ± 0.038 | 0.166 ± 0.017 | 0.354 ± 0.005 | 0.182 ± 0.013 |
| **Random Forest** |  |  |  |  |  |  |  |  |
| MSE | 2769.314 ± 63.752 | 2300.937 ± 41.956 | 2256.726 ± 57.050 | 2264.232 ± 64.790 | 2263.339 ± 61.481 | 2259.029 ± 50.935 | **2249.802 ± 52.171** | 2254.751 ± 49.772 |
| RMSE | 52.621 ± 0.607 | 47.966 ± 0.436 | 47.502 ± 0.597 | 47.580 ± 0.685 | 47.571 ± 0.649 | 47.527 ± 0.534 | **47.429 ± 0.550** | 47.482 ± 0.524 |
| MAE | 36.002 ± 0.164 | 32.883 ± 0.160 | 32.660 ± 0.182 | 32.666 ± 0.255 | 32.706 ± 0.268 | 32.654 ± 0.196 | **32.613 ± 0.226** | 32.627 ± 0.225 |
| $R^2$ | 0.360 ± 0.011 | 0.468 ± 0.009 | 0.479 ± 0.010 | 0.477 ± 0.011 | 0.477 ± 0.008 | 0.478 ± 0.010 | **0.480 ± 0.006** | 0.479 ± 0.009 |
| Pearson | 0.600 ± 0.009 | 0.685 ± 0.007 | 0.694 ± 0.007 | 0.692 ± 0.008 | 0.692 ± 0.006 | 0.693 ± 0.008 | **0.695 ± 0.004** | 0.694 ± 0.007 |
| Spearman | 0.511 ± 0.004 | 0.601 ± 0.005 | 0.603 ± 0.004 | 0.603 ± 0.004 | 0.603 ± 0.004 | 0.603 ± 0.004 | **0.604 ± 0.003** | **0.604 ± 0.005** |
| **XGBoost** |  |  |  |  |  |  |  |  |
| MSE | 2897.209 ± 43.137 | 2705.913 ± 49.775 | **2691.162 ± 43.130** | 2705.938 ± 53.598 | 2699.473 ± 56.277 | 2700.688 ± 51.372 | 2693.513 ± 58.623 | 2694.935 ± 55.838 |
| RMSE | 53.824 ± 0.400 | 52.016 ± 0.479 | **51.875 ± 0.416** | 52.016 ± 0.514 | 51.954 ± 0.541 | 51.966 ± 0.494 | 51.896 ± 0.561 | 51.910 ± 0.537 |
| MAE | 36.681 ± 0.222 | 35.704 ± 0.165 | 35.700 ± 0.170 | 35.759 ± 0.176 | 35.707 ± 0.225 | 35.730 ± 0.222 | 35.729 ± 0.187 | **35.693 ± 0.216** |
| $R^2$ | 0.330 ± 0.012 | 0.375 ± 0.006 | **0.378 ± 0.009** | 0.375 ± 0.009 | 0.376 ± 0.010 | 0.376 ± 0.009 | 0.378 ± 0.007 | 0.377 ± 0.011 |
| Pearson | 0.580 ± 0.012 | 0.629 ± 0.006 | **0.632 ± 0.009** | 0.629 ± 0.009 | 0.630 ± 0.009 | 0.630 ± 0.010 | **0.632 ± 0.007** | 0.631 ± 0.011 |
| Spearman | 0.478 ± 0.005 | 0.515 ± 0.004 | **0.514 ± 0.005** | 0.513 ± 0.006 | **0.514 ± 0.007** | 0.513 ± 0.005 | 0.513 ± 0.004 | **0.514 ± 0.005** |
| **SNN** |  |  |  |  |  |  |  |  |
| MSE | 2951.707 ± 48.938 | 3986.566 ± 122.736 | **2524.230 ± 73.636** | 3734.008 ± 87.264 | 3990.670 ± 72.496 | 4001.180 ± 109.496 | 3558.211 ± 71.228 | 3998.045 ± 102.851 |
| RMSE | 54.328 ± 0.452 | 63.133 ± 0.971 | **50.237 ± 0.733** | 61.103 ± 0.710 | 63.169 ± 0.572 | 63.250 ± 0.867 | 59.648 ± 0.597 | 63.225 ± 0.813 |
| MAE | 36.698 ± 0.198 | 41.079 ± 0.343 | **34.486 ± 0.249** | 40.292 ± 0.396 | 41.057 ± 0.226 | 41.088 ± 0.184 | 39.613 ± 0.250 | 41.096 ± 0.263 |
| $R^2$ | 0.318 ± 0.006 | 0.079 ± 0.004 | **0.417 ± 0.011** | 0.137 ± 0.010 | 0.078 ± 0.003 | 0.076 ± 0.005 | 0.178 ± 0.008 | 0.076 ± 0.004 |
| Pearson | 0.576 ± 0.006 | 0.298 ± 0.008 | **0.667 ± 0.009** | 0.417 ± 0.017 | 0.297 ± 0.007 | 0.295 ± 0.007 | 0.466 ± 0.013 | 0.294 ± 0.006 |
| Spearman | 0.486 ± 0.003 | 0.323 ± 0.008 | **0.555 ± 0.005** | 0.389 ± 0.006 | 0.322 ± 0.007 | 0.321 ± 0.006 | 0.413 ± 0.005 | 0.320 ± 0.007 |
| **SYNDEEP** |  |  |  |  |  |  |  |  |
| MSE | 21455.086 ± 19473.376 | 3786.669 ± 125.663 | **2661.424 ± 76.165** | 3580.770 ± 80.007 | 3792.805 ± 113.310 | 3801.472 ± 126.567 | 3547.407 ± 219.576 | 3790.025 ± 93.000 |
| RMSE | 132.358 ± 66.135 | 61.528 ± 1.018 | **51.584 ± 0.741** | 59.836 ± 0.670 | 61.580 ± 0.924 | 61.648 ± 1.024 | 59.535 ± 1.805 | 61.559 ± 0.759 |
| MAE | 39.375 ± 1.915 | 40.160 ± 0.253 | **35.233 ± 0.281** | 39.255 ± 0.290 | 40.170 ± 0.224 | 40.166 ± 0.339 | 39.040 ± 0.265 | 40.166 ± 0.301 |
| $R^2$ | -3.992 ± 4.572 | 0.125 ± 0.004 | **0.385 ± 0.017** | 0.173 ± 0.012 | 0.124 ± 0.005 | 0.122 ± 0.009 | 0.181 ± 0.031 | 0.124 ± 0.003 |
| Pearson | 0.326 ± 0.107 | 0.368 ± 0.007 | **0.663 ± 0.011** | 0.446 ± 0.016 | 0.366 ± 0.008 | 0.363 ± 0.019 | 0.450 ± 0.027 | 0.368 ± 0.005 |
| Spearman | 0.494 ± 0.003 | 0.361 ± 0.004 | **0.557 ± 0.004** | 0.409 ± 0.006 | 0.359 ± 0.007 | 0.361 ± 0.006 | 0.412 ± 0.005 | 0.361 ± 0.005 |
| **PCNNGL** |  |  |  |  |  |  |  |  |
| MSE | 3972.745 ± 582.792 | 3854.359 ± 209.849 | 3719.423 ± 452.836 | 3857.975 ± 325.480 | 3595.023 ± 178.989 | **3434.672 ± 198.031** | 3544.200 ± 190.825 | 3581.009 ± 520.798 |
| RMSE | 62.886 ± 4.491 | 62.063 ± 1.679 | 60.887 ± 3.682 | 62.063 ± 2.604 | 59.942 ± 1.500 | **58.585 ± 1.663** | 59.514 ± 1.589 | 59.718 ± 4.054 |
| MAE | 39.777 ± 0.458 | 39.495 ± 0.200 | 39.281 ± 0.571 | 39.427 ± 0.212 | 39.160 ± 0.268 | **38.949 ± 0.302** | 39.205 ± 0.411 | 39.061 ± 0.527 |
| $R^2$ | 0.082 ± 0.137 | 0.108 ± 0.061 | 0.142 ± 0.090 | 0.109 ± 0.074 | 0.169 ± 0.046 | **0.206 ± 0.050** | 0.181 ± 0.043 | 0.172 ± 0.121 |
| Pearson | 0.455 ± 0.020 | 0.477 ± 0.023 | 0.480 ± 0.030 | 0.475 ± 0.022 | 0.496 ± 0.021 | **0.503 ± 0.023** | 0.493 ± 0.015 | 0.499 ± 0.033 |
| Spearman | 0.404 ± 0.005 | 0.429 ± 0.005 | 0.429 ± 0.009 | 0.431 ± 0.003 | 0.433 ± 0.005 | **0.434 ± 0.005** | 0.430 ± 0.004 | 0.437 ± 0.004 |
| **PCNNGL-SYNDEEP** |  |  |  |  |  |  |  |  |
| MSE | 4313.141 ± 1429.700 | 4038.124 ± 94.647 | 4090.957 ± 1536.015 | 3986.251 ± 835.574 | 4039.243 ± 93.556 | 4032.705 ± 159.938 | **3627.088 ± 82.883** | 4028.255 ± 122.236 |
| RMSE | 64.997 ± 9.918 | 63.542 ± 0.745 | 63.236 ± 10.119 | 62.863 ± 6.196 | 63.551 ± 0.737 | 63.492 ± 1.262 | **60.222 ± 0.689** | 63.462 ± 0.966 |
| MAE | 40.027 ± 0.368 | 41.378 ± 0.279 | **39.841 ± 0.329** | 39.929 ± 0.398 | 41.386 ± 0.242 | 41.365 ± 0.348 | 39.903 ± 0.307 | 41.328 ± 0.253 |
| $R^2$ | -0.001 ± 0.352 | 0.067 ± 0.003 | 0.046 ± 0.402 | 0.078 ± 0.201 | 0.067 ± 0.003 | 0.068 ± 0.004 | **0.162 ± 0.015** | 0.069 ± 0.004 |
| Pearson | 0.406 ± 0.092 | 0.339 ± 0.011 | 0.427 ± 0.078 | 0.406 ± 0.078 | 0.338 ± 0.011 | 0.339 ± 0.009 | **0.459 ± 0.024** | 0.339 ± 0.010 |
| Spearman | 0.390 ± 0.007 | 0.333 ± 0.008 | **0.395 ± 0.005** | 0.392 ± 0.004 | 0.332 ± 0.008 | 0.333 ± 0.008 | **0.395 ± 0.005** | 0.331 ± 0.008 |

**Supplemental Table 5:** Mean and standard deviation across 10-fold cross validation performance for seven different machine learning models on the dose-dependent percent growth regression task. Each section corresponds to one of the following machine learning models including: support vector machine (SVM), random forest, eXtreme gradient boosting (XGBoost), shallow neural network (SNN), SYNDEEP [14], partially connected neural network with hidden gene layer (PCNNGL), and the PCNNGL-SYNDEEP hybrid model. The columns denote different input feature combinations: Morgan fingerprints (MF), MF with DNA, MF with RNA, MF with protein expression (PE), MF with DNA and RNA, MF with DNA and PE, MF with RNA and PE, and MF with all cell line feature modalities. Metrics in bold indicate the best performance across all input feature combinations for that model.

| Metric | MF | MF+DNA | MF+RNA | MF+PE | MF+DNA+RNA | MF+DNA+PE | MF+RNA+PE | MF+All |
| --- | --- | --- | --- | --- | --- | --- | --- | --- |
| **SVM** |  |  |  |  |  |  |  |  |
| MSE | 1905.853 ± 7.750 | 4602.569 ± 2836.581 | **1795.540 ± 10.431** | 1846.090 ± 174.427 | 3457.996 ± 1062.582 | 4404.918 ± 2513.912 | 1932.483 ± 127.687 | 3241.835 ± 793.875 |
| RMSE | 43.656 ± 0.089 | 65.165 ± 19.892 | **42.374 ± 0.123** | 42.925 ± 1.976 | 58.220 ± 8.718 | 64.327 ± 17.224 | 43.939 ± 1.435 | 56.584 ± 6.676 |
| MAE | 27.705 ± 0.056 | 53.030 ± 19.793 | **27.376 ± 0.080** | 28.413 ± 1.280 | 45.939 ± 9.816 | 52.210 ± 17.301 | 30.102 ± 0.669 | 43.866 ± 8.174 |
| $R^2$ | 0.068 ± 0.003 | -1.250 ± 1.388 | **0.122 ± 0.002** | 0.098 ± 0.085 | -0.690 ± 0.519 | -1.154 ± 1.233 | 0.055 ± 0.062 | -0.585 ± 0.390 |
| Pearson | 0.375 ± 0.002 | 0.296 ± 0.048 | **0.432 ± 0.002** | 0.421 ± 0.001 | 0.240 ± 0.035 | 0.240 ± 0.032 | 0.338 ± 0.006 | 0.258 ± 0.021 |
| Spearman | 0.445 ± 0.002 | 0.278 ± 0.036 | **0.466 ± 0.001** | 0.462 ± 0.001 | 0.222 ± 0.020 | 0.221 ± 0.018 | 0.349 ± 0.003 | 0.234 ± 0.022 |
| **Random Forest GPU** |  |  |  |  |  |  |  |  |
| MSE | 1337.693 ± 3.662 | 1307.500 ± 7.131 | 1294.645 ± 6.403 | 1293.546 ± 7.237 | 1294.010 ± 5.983 | **1289.810 ± 6.216** | 1294.341 ± 5.058 | 1294.600 ± 6.318 |
| RMSE | 36.574 ± 0.050 | 36.159 ± 0.099 | 35.981 ± 0.089 | 35.966 ± 0.101 | 35.972 ± 0.083 | **35.914 ± 0.086** | 35.977 ± 0.070 | 35.980 ± 0.088 |
| MAE | 25.947 ± 0.024 | **25.912 ± 0.048** | 26.367 ± 0.061 | 26.367 ± 0.059 | 26.362 ± 0.052 | 26.363 ± 0.050 | 26.362 ± 0.042 | 26.367 ± 0.053 |
| $R^2$ | 0.346 ± 0.002 | 0.361 ± 0.003 | 0.367 ± 0.001 | 0.368 ± 0.002 | 0.367 ± 0.001 | **0.370 ± 0.001** | 0.367 ± 0.002 | 0.367 ± 0.001 |
| Pearson | 0.597 ± 0.002 | 0.612 ± 0.003 | 0.615 ± 0.001 | 0.615 ± 0.002 | 0.615 ± 0.001 | **0.618 ± 0.001** | 0.615 ± 0.002 | 0.614 ± 0.001 |
| Spearman | **0.497 ± 0.003** | 0.482 ± 0.003 | 0.453 ± 0.001 | 0.449 ± 0.002 | 0.450 ± 0.002 | 0.452 ± 0.001 | 0.453 ± 0.002 | 0.450 ± 0.001 |
| **XGBoost** |  |  |  |  |  |  |  |  |
| MSE | 769.808 ± 3.140 | 464.616 ± 6.162 | 454.938 ± 4.148 | 459.302 ± 5.005 | 453.168 ± 3.221 | 455.000 ± 2.678 | **452.526 ± 3.126** | 454.661 ± 3.338 |
| RMSE | 27.745 ± 0.057 | 21.555 ± 0.143 | 21.329 ± 0.097 | 21.431 ± 0.117 | 21.288 ± 0.076 | 21.331 ± 0.063 | **21.273 ± 0.073** | 21.323 ± 0.078 |
| MAE | 17.699 ± 0.033 | 14.794 ± 0.092 | 14.773 ± 0.063 | 14.839 ± 0.078 | 14.736 ± 0.049 | 14.763 ± 0.031 | **14.723 ± 0.038** | 14.749 ± 0.060 |
| $R^2$ | 0.624 ± 0.002 | 0.773 ± 0.002 | 0.778 ± 0.002 | 0.775 ± 0.002 | 0.778 ± 0.002 | 0.778 ± 0.002 | **0.779 ± 0.001** | 0.778 ± 0.002 |
| Pearson | 0.791 ± 0.001 | 0.884 ± 0.001 | 0.886 ± 0.001 | 0.885 ± 0.001 | **0.887 ± 0.001** | 0.886 ± 0.001 | **0.887 ± 0.001** | 0.886 ± 0.001 |
| Spearman | 0.747 ± 0.001 | 0.775 ± 0.002 | **0.776 ± 0.001** | 0.775 ± 0.001 | **0.776 ± 0.001** | 0.775 ± 0.002 | **0.776 ± 0.001** | **0.776 ± 0.001** |
| **SNN** |  |  |  |  |  |  |  |  |
| MSE | 1602.257 ± 5.308 | 1575.714 ± 7.417 | **1347.958 ± 4.051** | 1534.020 ± 10.070 | 1567.512 ± 6.076 | 1572.099 ± 7.979 | 1513.827 ± 7.822 | 1573.069 ± 7.135 |
| RMSE | 40.028 ± 0.066 | 39.695 ± 0.093 | **36.715 ± 0.055** | 39.166 ± 0.129 | 39.592 ± 0.077 | 39.650 ± 0.101 | 38.908 ± 0.101 | 39.662 ± 0.090 |
| MAE | 28.806 ± 0.082 | 29.242 ± 0.363 | **26.867 ± 0.078** | 28.629 ± 0.282 | 28.829 ± 0.321 | 29.057 ± 0.312 | 28.471 ± 0.146 | 29.068 ± 0.294 |
| $R^2$ | 0.217 ± 0.001 | 0.230 ± 0.003 | **0.341 ± 0.001** | 0.250 ± 0.003 | 0.234 ± 0.002 | 0.232 ± 0.003 | 0.260 ± 0.004 | 0.231 ± 0.003 |
| Pearson | 0.469 ± 0.002 | 0.493 ± 0.002 | **0.590 ± 0.001** | 0.514 ± 0.003 | 0.494 ± 0.002 | 0.494 ± 0.001 | 0.523 ± 0.005 | 0.493 ± 0.003 |
| Spearman | 0.479 ± 0.002 | 0.479 ± 0.002 | **0.544 ± 0.002** | 0.498 ± 0.001 | 0.478 ± 0.002 | 0.479 ± 0.002 | 0.504 ± 0.003 | 0.478 ± 0.003 |
| **SYNDEEP** |  |  |  |  |  |  |  |  |
| MSE | 1639.362 ± 69.939 | 1506.756 ± 5.833 | **1328.185 ± 3.213** | 1488.480 ± 10.605 | 1506.341 ± 7.596 | 1506.204 ± 10.023 | 1463.035 ± 9.133 | 1504.383 ± 13.037 |
| RMSE | 40.481 ± 0.853 | 38.817 ± 0.075 | **36.444 ± 0.044** | 38.581 ± 0.137 | 38.811 ± 0.098 | 38.810 ± 0.129 | 38.249 ± 0.119 | 38.786 ± 0.168 |
| MAE | 28.838 ± 0.192 | 28.716 ± 0.106 | **26.639 ± 0.087** | 28.494 ± 0.162 | 28.695 ± 0.156 | 28.698 ± 0.152 | 28.171 ± 0.147 | 28.665 ± 0.170 |
| $R^2$ | 0.199 ± 0.034 | 0.263 ± 0.003 | **0.351 ± 0.003** | 0.272 ± 0.006 | 0.264 ± 0.003 | 0.264 ± 0.003 | 0.285 ± 0.004 | 0.265 ± 0.004 |
| Pearson | 0.452 ± 0.034 | 0.524 ± 0.002 | **0.604 ± 0.002** | 0.538 ± 0.004 | 0.524 ± 0.003 | 0.524 ± 0.003 | 0.547 ± 0.003 | 0.524 ± 0.003 |
| Spearman | 0.486 ± 0.001 | 0.474 ± 0.003 | **0.557 ± 0.002** | 0.494 ± 0.006 | 0.473 ± 0.004 | 0.474 ± 0.003 | 0.500 ± 0.005 | 0.474 ± 0.003 |
| **PCNNGL** |  |  |  |  |  |  |  |  |
| MSE | 1455.367 ± 163.421 | 1442.625 ± 299.786 | 1348.187 ± 279.181 | 1313.749 ± 204.091 | 1290.198 ± 179.755 | 1314.238 ± 108.497 | 1292.485 ± 225.601 | **1224.326 ± 45.269** |
| RMSE | 38.100 ± 2.051 | 37.813 ± 3.772 | 36.565 ± 3.524 | 36.160 ± 2.624 | 35.850 ± 2.352 | 36.225 ± 1.482 | 35.849 ± 2.861 | **34.985 ± 0.639** |
| MAE | 25.864 ± 0.564 | 25.424 ± 0.735 | **24.864 ± 0.410** | 25.042 ± 0.501 | 25.030 ± 0.362 | 25.363 ± 0.571 | 25.076 ± 0.480 | 25.038 ± 0.913 |
| $R^2$ | 0.289 ± 0.080 | 0.295 ± 0.147 | 0.341 ± 0.134 | 0.358 ± 0.099 | 0.369 ± 0.090 | 0.358 ± 0.051 | 0.368 ± 0.112 | **0.402 ± 0.020** |
| Pearson | 0.551 ± 0.047 | 0.580 ± 0.070 | 0.605 ± 0.066 | 0.611 ± 0.054 | 0.620 ± 0.045 | 0.609 ± 0.033 | 0.622 ± 0.053 | **0.639 ± 0.013** |
| Spearman | 0.569 ± 0.003 | 0.578 ± 0.004 | **0.582 ± 0.003** | 0.578 ± 0.004 | **0.582 ± 0.003** | 0.580 ± 0.003 | **0.582 ± 0.004** | **0.582 ± 0.007** |
| **PCNNGL-SYNDEEP** |  |  |  |  |  |  |  |  |
| MSE | 1687.378 ± 10.325 | 1705.254 ± 10.608 | **1661.310 ± 10.025** | 1684.865 ± 3.397 | 1705.075 ± 10.625 | 1708.252 ± 9.840 | 1662.217 ± 7.714 | 1705.494 ± 9.999 |
| RMSE | 41.078 ± 0.126 | 41.295 ± 0.128 | **40.759 ± 0.123** | 41.047 ± 0.041 | 41.292 ± 0.129 | 41.331 ± 0.119 | 40.770 ± 0.095 | 41.297 ± 0.121 |
| MAE | 30.527 ± 0.206 | 31.692 ± 0.135 | **30.104 ± 0.213** | 30.409 ± 0.163 | 31.628 ± 0.192 | 31.726 ± 0.054 | 30.177 ± 0.139 | 31.719 ± 0.176 |
| $R^2$ | 0.175 ± 0.002 | 0.166 ± 0.003 | **0.188 ± 0.002** | 0.176 ± 0.001 | 0.167 ± 0.004 | 0.165 ± 0.002 | **0.188 ± 0.001** | 0.166 ± 0.003 |
| Pearson | 0.438 ± 0.002 | 0.442 ± 0.003 | **0.458 ± 0.003** | 0.440 ± 0.002 | 0.441 ± 0.005 | 0.441 ± 0.002 | 0.457 ± 0.002 | 0.441 ± 0.004 |
| Spearman | 0.448 ± 0.002 | 0.386 ± 0.005 | **0.457 ± 0.002** | 0.449 ± 0.002 | 0.386 ± 0.007 | 0.387 ± 0.004 | **0.457 ± 0.001** | 0.385 ± 0.005 |

**Supplemental Figure 1:**
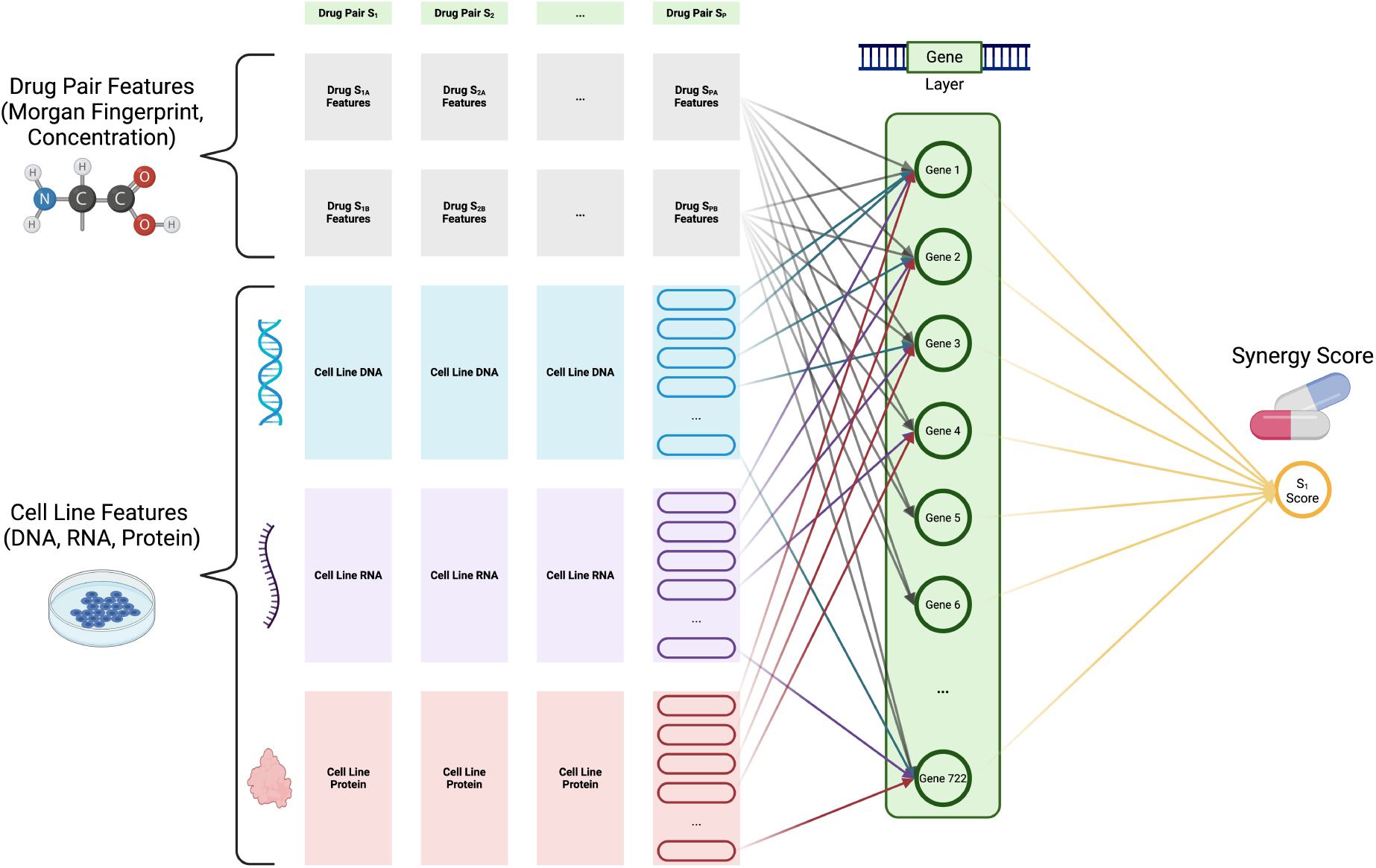
Diagram displaying architecture of partially connected neural network with hidden gene layer (PCNNGL) model. The PCNNGL model contains one hidden layer where nodes correspond to specific genes before feeding into the output layer. The top rows of gray boxes displays examples of drug pair features. Each column represents a single training data point, containing drug pair and cell line features. Each drug *A* and *B* has both Morgan fingerprint structure representations and concentration information, which are fully connected to every node in the hidden layer. The colored boxes represent cell line features of DNA, RNA, and protein expression. These cell line features are only connected to a hidden layer node if that data corresponds to the same gene, creating partial connections. All gene nodes in the hidden layer are fully connected to the output node.

**Supplemental Figure 2:**
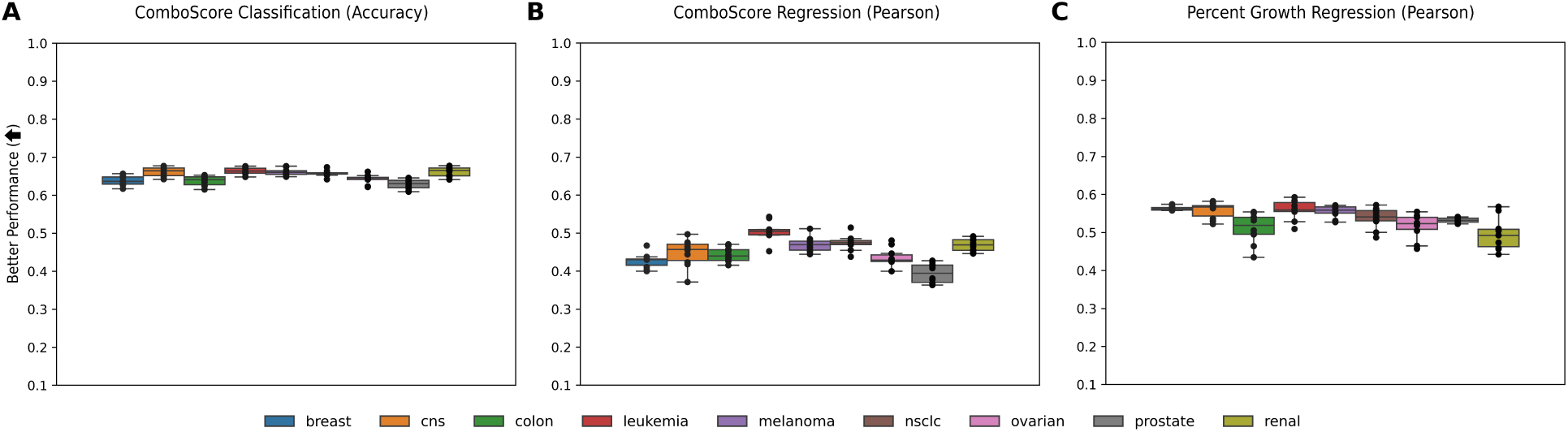
PCNNGL 10-fold cross validation model performance on individual tissue types. Tissue types include breast, central nervous system (CNS), colon, leukemia, melanoma, non-small lung cell carcinoma (NSCLC), ovarian, prostate, and renal. Box and whisker plots of the 10-fold cross validation performance are given for **(A)** accuracy on ComboScore classification task, **(B)** Pearson correlation on the ComboScore regression task, and **(C)** Pearson correlation on the percent growth regression task.

**Supplemental Figure 3:**
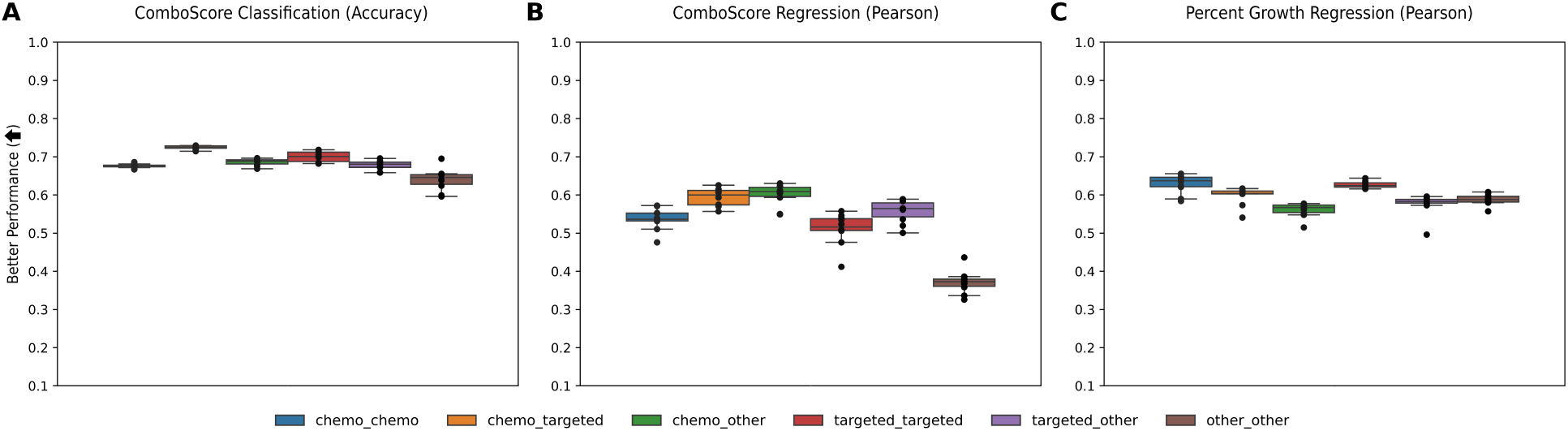
PCNNGL 10-fold cross validation model performance on specific drug classes. Classes of drug combinations include: chemotherapy-chemotherapy (chemo_chemo), chemotherapy-targeted therapy (chemo_targeted), chemotherapy-other (chemo_other), targeted therapy-targeted therapy (targeted_targeted), targeted therapy-other (targeted_other), and other-other (other_other). Box and whisker plots of the 10-fold cross validation performance are given for **(A)** accuracy on ComboScore classification task, **(B)** Pearson correlation on the ComboScore regression task, and **(C)** Pearson correlation on the percent growth regression task.

**Supplemental Figure 4:**
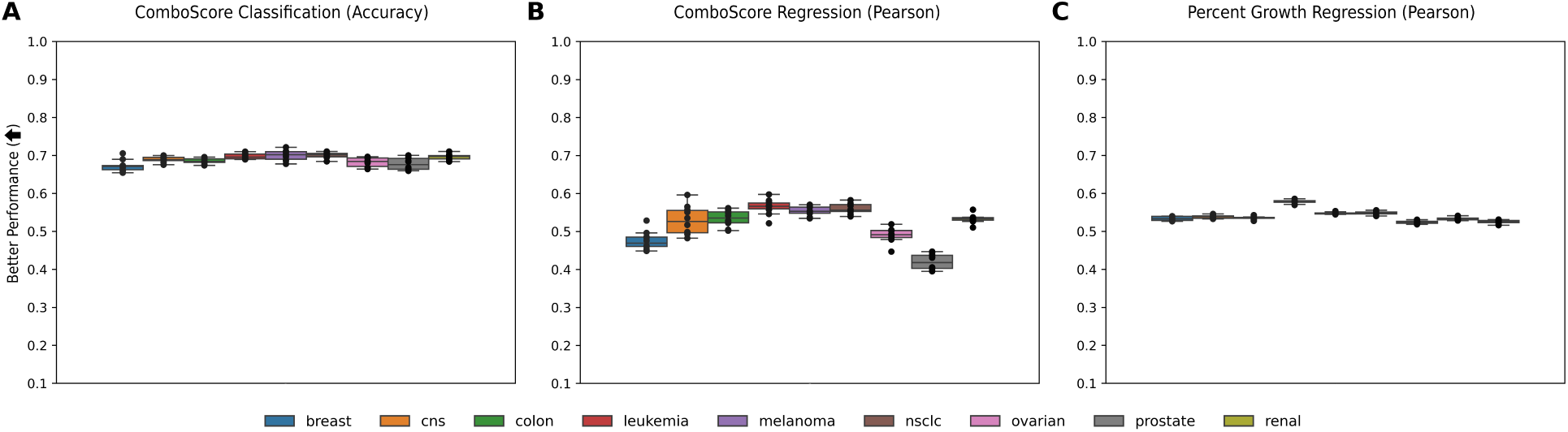
SNN 10-fold cross validation model performance on individual tissue types. Tissue types include breast, central nervous system (CNS), colon, leukemia, melanoma, non-small lung cell carcinoma (NSCLC), ovarian, prostate, and renal. Box and whisker plots of the 10-fold cross validation performance are given for **(A)** accuracy on ComboScore classification task, **(B)** Pearson correlation on the ComboScore regression task, and **(C)** Pearson correlation on the percent growth regression task.

**Supplemental Figure 5:**
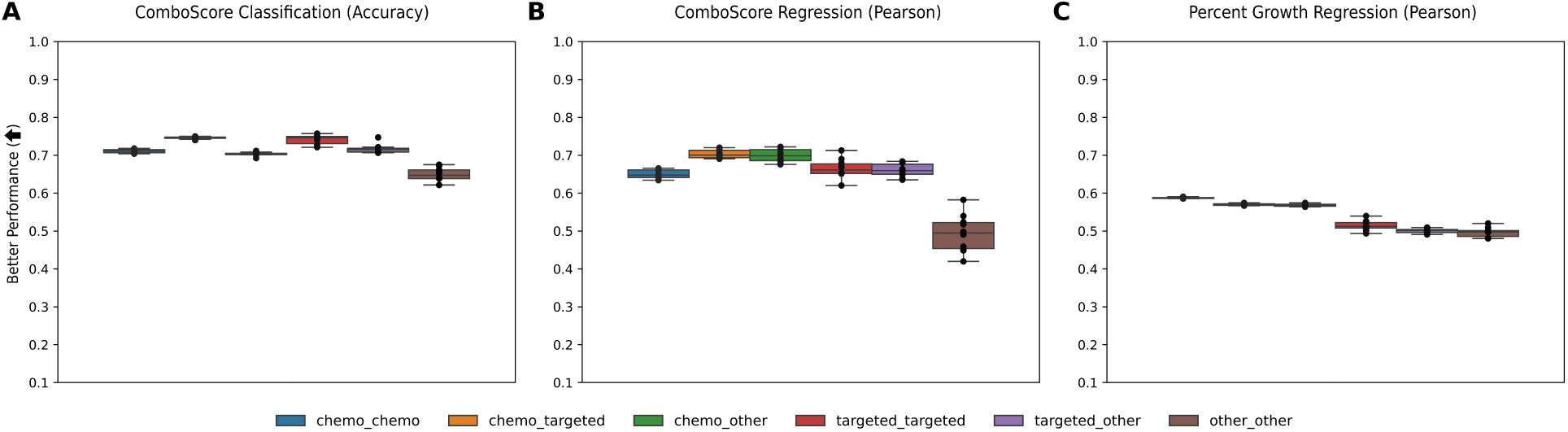
SNN 10-fold cross validation model performance on specific drug classes. Classes of drug combinations include: chemotherapy-chemotherapy (chemo_chemo), chemotherapy-targeted therapy (chemo_targeted), chemotherapy-other (chemo_other), targeted therapy-targeted therapy (targeted_targeted), targeted therapy-other (targeted_other), and other-other (other_other). Box and whisker plots of the 10-fold cross validation performance are given for **(A)** accuracy on ComboScore classification task, **(B)** Pearson correlation on the ComboScore regression task, and **(C)** Pearson correlation on the percent growth regression task.

**Supplemental Figure 6:**
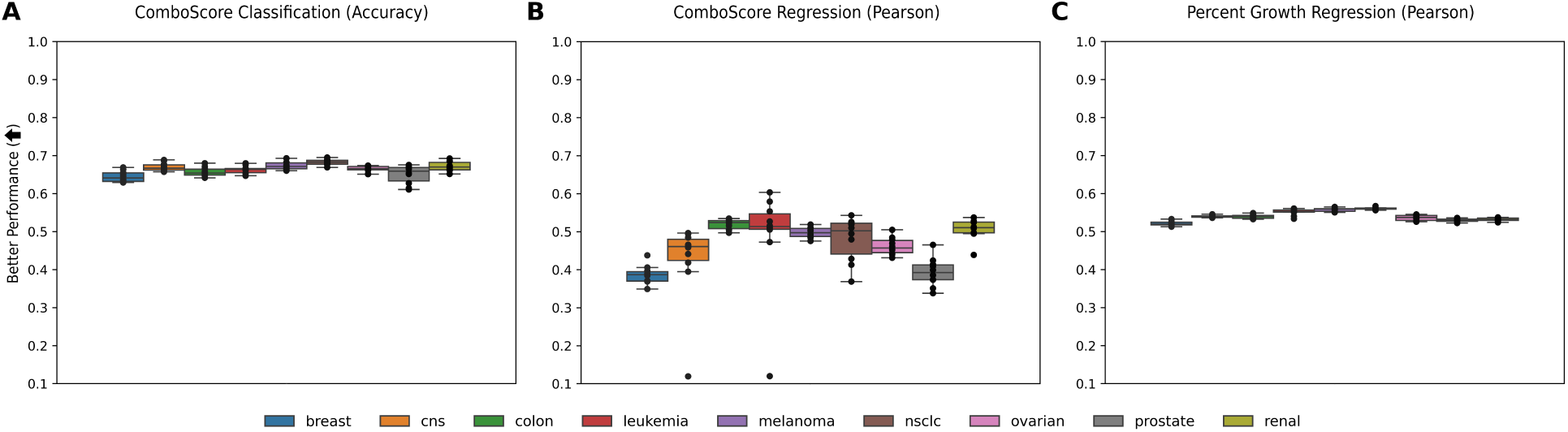
SYNDEEP 10-fold cross validation model performance on individual tissue types. Tissue types include breast, central nervous system (CNS), colon, leukemia, melanoma, non-small lung cell carcinoma (NSCLC), ovarian, prostate, and renal. Box and whisker plots of the 10-fold cross validation performance are given for **(A)** accuracy on ComboScore classification task, **(B)** Pearson correlation on the ComboScore regression task, and **(C)** Pearson correlation on the percent growth regression task.

**Supplemental Figure 7:**
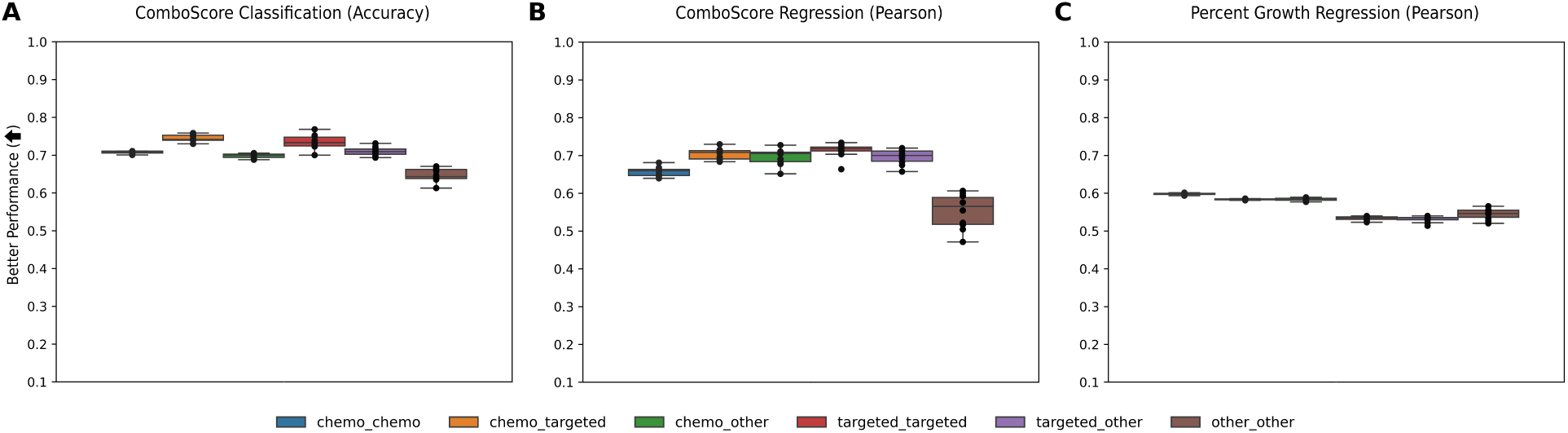
SYNDEEP 10-fold cross validation model performance on specific drug classes. Classes of drug combinations include: chemotherapy-chemotherapy (chemo_chemo), chemotherapy-targeted therapy (chemo_targeted), chemotherapy-other (chemo_other), targeted therapy-targeted therapy (targeted_targeted), targeted therapy-other (targeted_other), and other-other (other_other). Box and whisker plots of the 10-fold cross validation performance are given for **(A)** accuracy on ComboScore classification task, **(B)** Pearson correlation on the ComboScore regression task, and **(C)** Pearson correlation on the percent growth regression task.

**Supplemental Figure 8:**
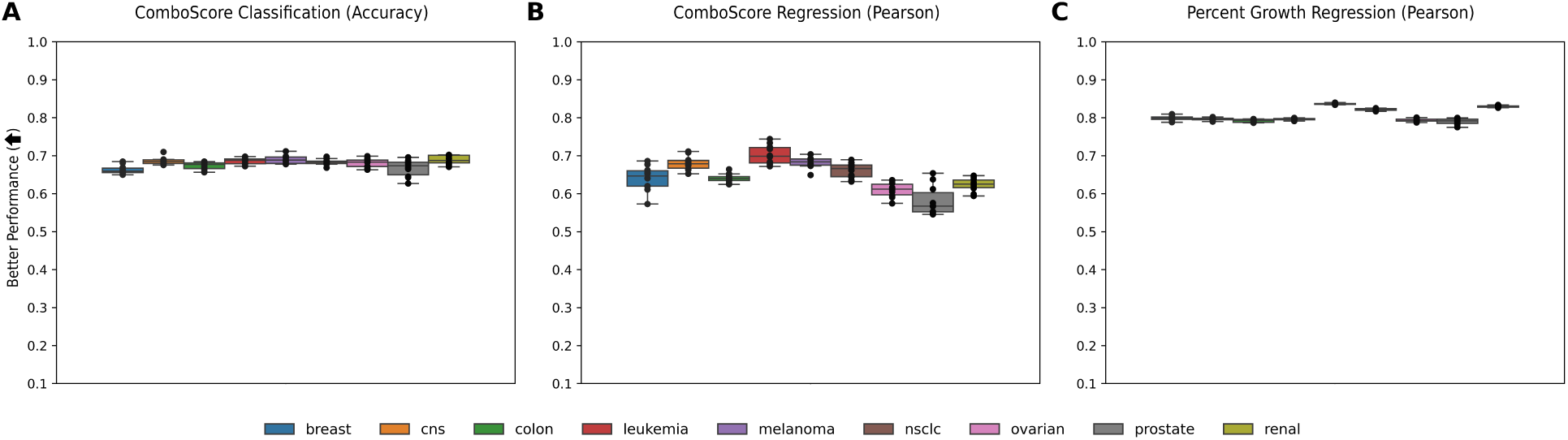
RF 10-fold cross validation model performance on individual tissue types. Tissue types include breast, central nervous system (CNS), colon, leukemia, melanoma, non-small lung cell carcinoma (NSCLC), ovarian, prostate, and renal. Box and whisker plots of the 10-fold cross validation performance are given for **(A)** accuracy on ComboScore classification task, **(B)** Pearson correlation on the ComboScore regression task, and **(C)** Pearson correlation on the percent growth regression task.

**Supplemental Figure 9:**
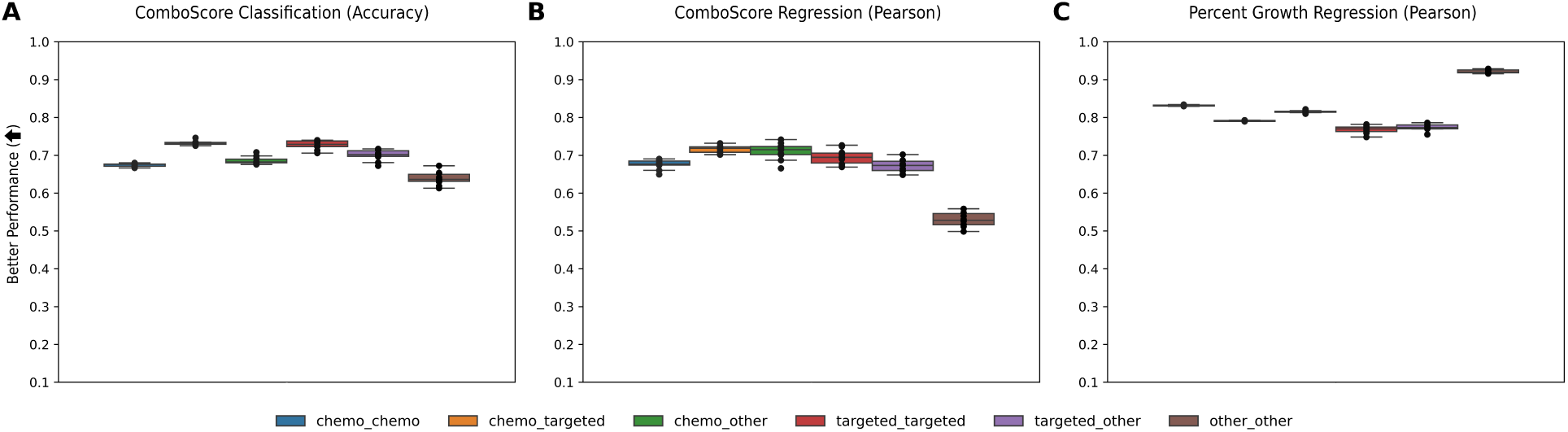
RF 10-fold cross validation model performance on specific drug classes. Classes of drug combinations include: chemotherapy-chemotherapy (chemo_chemo), chemotherapy-targeted therapy (chemo_targeted), chemotherapy-other (chemo_other), targeted therapy-targeted therapy (targeted_targeted), targeted therapy-other (targeted_other), and other-other (other_other). Box and whisker plots of the 10-fold cross validation performance are given for **(A)** accuracy on ComboScore classification task, **(B)** Pearson correlation on the ComboScore regression task, and **(C)** Pearson correlation on the percent growth regression task.

## Footnotes

1 Due to limited computing resources, RF cannot be run on the entire percent growth dataset using scikit-learn and a CPU. The ComboScore dataset only includes 300,928 samples because it summarizes over concentrations; however, the percent growth data set has 2,774,280 samples because it is dose-dependent. As a result, for the percent growth data, we implement a GPU compatible RF using the XGBRFRegressor function from the Chen and Guestrin [43] software package with the same hyper-parameter settings. Note that these implementations of the algorithm are slightly different (e.g., see Supplementary Tables 1 and 2, respectively). As such, we suspect that the performance of the GPU RF algorithm from XGBoost maybe less optimized than the scikit-learn implementation.

